# Deep evolutionary origins of the connexin gene family

**DOI:** 10.64898/2026.06.13.731801

**Authors:** Jack Butler, Julia Aniol, Nadine Hillman, Sarbjit Nijjar, Nicholas Dale, Edward G Smith

## Abstract

The vertebrate connexin family encodes a diverse repertoire of transmembrane channels that mediate both intercellular communication and cytosol to extracellular space exchange. Their apparent absence in non-chordates has led to the prevailing view that connexins arose during chordate evolution with analogous functions performed in non-chordates by the innexins, a structurally similar but non-homologous protein family. Here, using a combination of sequence and structural analyses, we discover the presence of connexins in anthozoan and protostome genomes despite widespread secondary gene loss in Bilateria. We show that these connexins can be present in up to 100 copies per genome and are often found as N-terminal domains of NOD-like receptors (NLRs) or other immune genes. We show that these invertebrate connexins localise at the cell membrane, function as either hemichannels (anthozoans and protostomes) or gap junctions (protostomes), and respond to depolarisation enabling the transfer of small molecules. Further, we demonstrate that a subset of the invertebrate connexin-like domains are sensitive to CO_2_ and mirroring findings in vertebrate connexins, the CO_2_-mediated gating stems from the formation of a carbamate bridge between neighbouring subunits. Taken together, we trace the evolution of this gene family to the cnidarian-bilaterian ancestor, demonstrate functional similarity between invertebrate and vertebrate connexins, and suggest an ancient role of connexins in innate immunity.

## Introduction

Communication and coordination between cells are essential processes in multicellular life. In animals, electrical and chemical signalling between adjacent cells can be performed by large-pore plasma membrane channels ^1–4^ that form gap junction channels between adjacent cells. These transmembrane channel proteins belong to two non-homologous families, the connexins and the innexins. Gap junction proteins are found in nearly all metazoans and dysfunction of connexin-mediated communication is linked to conditions such as non-syndromic deafness ^5^, keratitis ichthyosis deafness syndrome ^6^, X-linked Charcot Marie Tooth disease ^7, 8^ and oculodentodigital dysplasia ^9^. Understanding the evolutionary origins of these protein families, and connexins in particular, is essential to understanding the evolution of intercellular signalling and the development of new treatments for aberrant connexin-mediated signalling.

The connexin gene family is often referred to as a chordate innovation ^10^. The gene encodes plasma membrane channels formed of six subunits that have a large pore permeable to small molecules. The canonical function of connexins is to form gap junction channels. Here the separate hexameric channels in neighbouring and closely apposed cells dock together to form a dodecameric assembly that permits diffusion of water, ions and solutes between cells. The hexameric connexin assemblies - so called hemichannels -also have biological functions and serve to allow release of small molecules such as ATP into the extracellular space where they may perform signalling functions ^11–14^. Normally hemichannels are shut under resting conditions but can be opened, to varying extents depending on the subtype, by physiological stimuli such as changes in transmembrane voltage ^15–18^, increases in intracellular Ca^2+^ concentration ^19^ and small elevations of PCO_2_ ^20–22^.

The human genome contains 21 distinct connexin genes and this large gene family has arisen through both whole genome duplication events and gene duplications ^23, 24, 25^. All vertebrates, from the most primitive extant representatives (e.g. lamprey) to mammals have multiple connexins genes in their genomes. The connexins in the lamprey genome have orthologues in other vertebrates and show similar syntenic organisation, but lampreys possess fewer genes in this family. Connexin genes are also found in non-vertebrate chordates such as the tunicates, but these do not directly correspond to the subfamilies structure observed in vertebrates, instead reflecting an independent radiation ^26, 27^. Tunicate connexin genes have a more varied gene structure with several being split into multiple exons ^26, 28^. By contrast, the coding region of mammalian connexins is almost always within a single exon ^28^.

The presence of several connexin genes in these ancient chordate groups *(Agnatha and Tunicata)* ^29–31^, suggests that the origins of the gene family (from a single ancestral gene) may have been within a chordate group that is no longer extant, or could have predated the evolution of the chordates. It is intriguing therefore that connexin domains have been annotated as occurring at the N-termini of Cnidarian Nod-like receptors ^32, 33^ (herein referred to as connexin-NLRs). Here, we establish that connexin-like (CxL) domains are present in invertebrate taxa and extend their origins to at least the cnidarian-bilaterian ancestor. Using sequence and structural approaches, we identify CxL domains from three invertebrate phyla and demonstrate that these domains are distinct from innexins and pannexins (innexin homologs found in vertebrates ^34^). We show that these domains form plasma membrane channels that are permeable to signalling molecules such as ATP and glutamate. Furthermore, we show that CO_2_ sensitivity, a distinct gating mechanism of a subset of chordate connexins, is also present in invertebrate CxLs. Taken together, our findings push the evolutionary origins of connexins domains beyond ∼600Mya, establish that key fundamental features of chordate connexins are retained in their distant relatives and indicate an ancestral role of connexins in innate immunity.

## Results

### Presence of connexin-like domains in non-chordate NLRs

Observations of connexin domains in anthozoan proteins counters the prevailing hypothesis that connexins are exclusive to chordates ^10, 27, 35^ and suggests there may be unrecognised diversity in the connexin family. To evaluate the prevalence of connexin domains among non-chordate taxa, we searched for proteins possessing members of the Connexin N-terminal domain superfamily (IPR038359), based on the CATH-Gene3D protein structural classification, in the Uniprot database. After filtering (see Methods), this screen revealed 170 non-chordate proteins containing the IRP038359 domain (Supplementary Table 1). Of these, 158 belonged to the subphylum anthozoa, with additional records belonging to the annelid *Owenia fusiformis* (10) and the brachiopod, *Lingula anatina* (2). Curiously, while anthozoans possess innexins, these putative connexins domain lacked an innexin annotation suggesting that they represent a distinct domain.

To further investigate whether these domains are indeed distinct from invertebrate innexins, we performed a network clustering analysis of Uniprot domains belonging to connexin (IPR038359), innexin (IPR000990), and pannexin (IPR039099) domain families (Fig 1A). The anthozoan domains annotated as part of the connexin superfamily represent a distinct cluster from invertebrate innexin domains (including anthozoan innexins/pannexins) and share significant links with chordate connexins (Fig 1A; Supplementary Fig 1). No innexin sequence formed a significant link to any CxL representative and even under a more permissive threshold only a single query (out of 210 query sequences) matches a pannexin domain compared to 159 matches to chordate connexin domains (Supplementary Fig 2), indicating that CxLs share detectable primary-sequence similarity with connexins but not with innexins.

**Figure 1.**
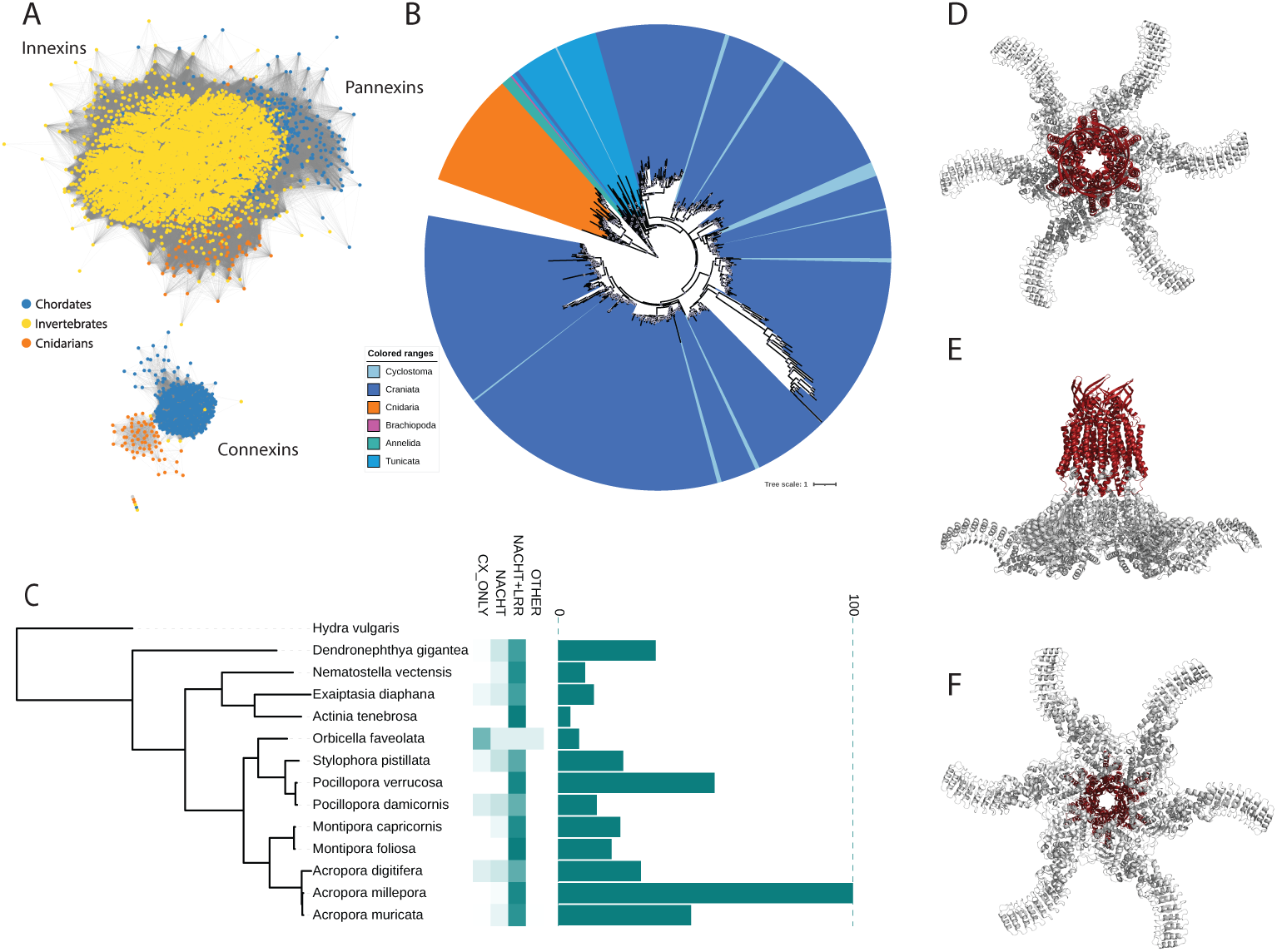
Non-chordate connexin domains are distinct from innexins and commonly found as NLRs. **A)** Clustering network based on sequence similarity of protein domains (UniRef50) annotated as members of the connexin (IPR038359), innexin (IPR000990), and pannexin (IPR039099) domain families. Nodes are coloured according to taxonomy (Invertebrates excluding cnidarians and tunicates – yellow; cnidarians – orange; chordates – dark blue). Edges connect nodes sharing significant similarity (e-value < 1e^-5^). **B)** Maximum likelihood tree of connexin and connexin-like domains shaded according to taxonomy. **C)** Connexin-like domains are abundant in anthozoans and predominantly associated with NACHT and LRR domains. Left: Phylogeny of sampled anthozoans based on single copy orthologs with the hydrozoan *Hydra vulgaris* as the outgroup. Middle: Heatmap based on the proportion of CxL-domain proteins that contain the NLR associated NACHT and LRR domains. Right: Number of copies of CxL-domain containing proteins within the respective genomes. **D-F)** Predicted structure of a CxL-NLR from the sea anemone *Nematostella vectensis*. The connexin domain is highlighted in red.

We next validated our findings using hidden-markov model (HMM) and structural approaches which are well-suited to remote homology applications ^36–38^. Firstly, we generated a profile HMM from a curated alignment of anthozoan CxLs (but excluding representatives with the PFAM PF00029 connexin annotation) and queried the profile against the PDB_mmCIF70 and PFAM databases using HHPred ^39^. The top hits support homology of CxLs with connexins (Supplementary Tables 2 and 3), revealing shared conservation of sequence signatures with Gap junction gamma-3 (6L3U_A; Probability 99.95%, E-value 4.8x10^-27^) and the connexin PFAM PF00029 (Probability 99.56%, E-value 4.7x10^-15^) HMMs. Next, we took representatives from anthozoan CxLs and innexins, and queried their predicted structures against the AFDB-PROTEOME using Foldseek ^38^. The top hits for the CxLs were consistently chordate connexins, similarly, the top hits for anthozoan innexins were innexins/pannexins (Supplementary Fig 3; Supplementary Table 4). It is worth noting that there are two cysteine residues in the extracellular loops of anthozoan CxLs which is a feature commonly associated with innexins ^40^, however, this signature is also present in vertebrate connexins belonging to the epsilon clade ^27^. Taken together, the annotation of these domains as part of the connexin superfamily is supported by complementary lines of evidence based on sequence clustering, conserved domain signatures, and structural similarity support the presence of CxL domains in some invertebrate lineages. These findings revise the evolutionary history of the connexins, suggesting the presence of CxL domains in the last common cnidarian-bilaterian ancestor.

Anthozoan connexin-like (CxL) domains form a distinct clade among connexin and connexin-like domains alongside protostome representatives and urochordate connexins (Fig 1B). Targeted surveys of cnidarian genomes revealed that CxL domains are widespread across anthozoans yet absent from non-anthozoan cnidarians (e.g., hydrozoans) (Fig 1C; Supplementary Table 5). In the genomes of corals, the number of copies of CxL domains ranged from 4 to 100 with evidence of species-specific expansions (Fig 1C; Supplementary Fig 4). These genomes also contain innexins, however, at a lower frequency (0-1 genes per genome) than their non-anthozoan counterparts (Supplementary Table 5). Conversely, innexins are abundant in the genomes of the connexin-hosting protostomes, *O. fusiformis*and *L. anatina* (Supplementary Table 5).

A notable feature of the anthozoan proteins with the connexin superfamily annotation is that the CxL domain is predominantly located in the N terminus alongside a NACHT and leucine rich repeat (LRR) domains. The presence of a central NACHT domain and C-terminal LRRs are defining features of Nucleotide-binding domain and Leucine-rich Repeat (NLR) genes, a family of pattern recognition receptors involved in innate immunity. The presumably membrane-bound nature of these CxL-NLRs is a rare feature as NLRs are characteristically cytosolic receptors^41^. Ninety two percent of anthozoan CxL proteins possess a NACHT domain (Fig 1C; Supplementary Fig 4) and these NACHT domains belong to a distinct clade compared to other anthozoan NLRs ^42^. The remaining anthozoan CxL proteins that lack the NACHT domain could be a result of incomplete annotations, however, these non-NLR connexins largely fall within a single clade (Supplementary Fig 4). Interestingly, the NLR members of this clade possess a distinct gene structure compared to other anthozoan CxL-NLRs genes with an exon boundary located at the end of the CxL-like domain; in contrast, CxL-NLRs are typically single exon genes. A consistent feature of the connexin-like domain is that the coding sequences are maintained as a single exon in the anthozoans and this is consistent with the largely single coding exon nature of mammalian connexins ^40^. Intriguingly, CxL domains in annelid *O. fusiformis* – associated with domains such as STING, RHIM, and P-loop NTPases - possess both single and multiple coding exon CxL domains (Supplementary Table 6) with the latter typical of tunicate, lamprey, and some fish connexins ^26, 28^.

### Connexin-like domains share structural and functional properties with chordate connexins

We selected CxL domain-containing proteins from our screening for functional characterisation (Table 1). The predicted fold of the connexin domain of a single connexin-NLR subunit is very similar to that of a single mammalian connexin subunit. As connexins are hexamers, we used AlphaFold3 to assemble the individual connexin-NLR subunits into hexamers (Fig 1D-F; Supplementary Fig 5). Just like a mammalian connexin, the connexin-NLR proteins were predicted to form a channel with a large pore, and the N-terminus projecting into the pore with the potential to act as a gate. Other features of the structure were similar to that of a connexin, notably the extracellular loops containing antiparallel beta sheets. To check whether different stoichiometries were plausible, we varied the number of subunits per channel from 4 to 8. Hexameric and heptameric predicted structures looked the most plausible with ipTM and pTM scores above 0.7 for many of the predicted structures (Supplementary Fig 6).

**Table 1.** The CxL molecules and connexins studied, with originating species, accession number, presence of NLR domains and carbamylation motif and taxonomic grouping. The name of each molecule is either according to the literature (for the connexins) or the naming convention described in the text.

| Name | Species | Accession | NACH<br>T | Carbam.<br>Motif | Phylum |
| --- | --- | --- | --- | --- | --- |
| pmCxL30 | *Pocillopora meandrina* | A0A3M6TUS9_PO<br>CDA | No | Yes | Cnidaria |
| pmCxL62 | *Pocillopora meandrina* | A0AAU9X471_9CN<br>ID | Yes | Yes | Cnidaria |
| plCxL60 | *Porites lobata* | CAH3037244.1 | No | Yes | Cnidaria |
| dpCxL52 | *Desmophyllum pertusum* | A0A9W9Y920_9CN<br>ID | No | Yes | Cnidaria |
| edCxL50 | *Exaiptasia diaphana* | A0A913YIK5_EXA<br>DI | Yes | No | Cnidaria |
| ofCxL26 | *Owenia fusiformis* | A0A8J1U2R9_OWE<br>FU | No | Yes | Annelid |
| ofCxL40 | *Owenia fusiformis* | A0A8J1TTA1_OWE<br>FU | No | No | Annelid |
| laCxL27 | *Lingula anatina* | XP_013391505.1 | No | No | Brachiopod |
| ciCx30.9 | *Ciona intestinalis* | F6T0Z3 | No | Yes | Chordate<br>(Tunicata) |
| Cx27.5 | *Petromyzon marinus* | S4RXA7 | No | Yes | Chordate<br>(Agnatha) |
| Cx46 | *Petromyzon marinus* | S4RAX6 | No | Yes | Chordate<br>(Agnatha) |

The N-terminal helices of many connexins project within the pore and contain charged residues. As these charged helices experience the transmembrane electric field, they impart voltage sensitivity to hemichannel gating. In the anthozoan CxL pmCxL30 (pm = *Pocillopora meandrina*; CxL connexin-like domain; 30 kDa), the N-terminal helix (_1_M**D**A**E**AL**KK**I_9_) has 4 charged residues suggesting that pmCxL30 and any other connexin-NLRs with similarly charged N-terminal helices could exhibit voltage dependent gating.

To test whether the Cnidarian CxL genes could indeed form membrane channels that are functionally similar to vertebrate connexins, we selected pmCxL30, tagged the gene with mCherry at the C-terminus and cloned it into a mammalian expression vector. When expressed in HeLa cells pmCxL30 was present in a punctate pattern reminiscent of that observed for connexins such as Cx26 or Cx43 with clear localisation at the plasma membrane (Fig 2A).

**Figure 2.**
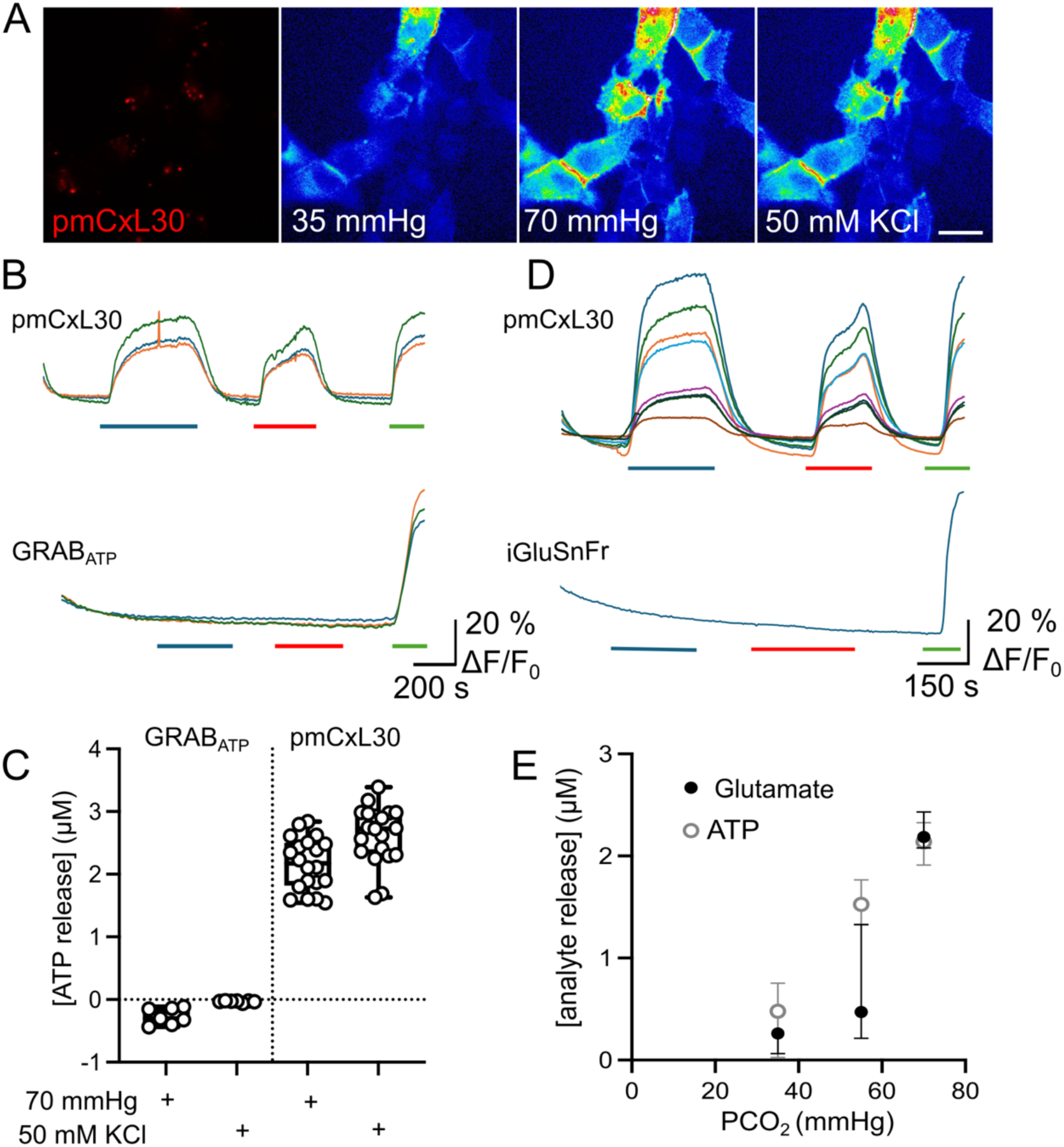
Cnidarian connexin-NLRs form connexin-like plasma membrane channels that can be gated by depolarisation and CO_2_. **A)** Images of expression of mCherry tagged pmCxL30 in HeLa cells (left) and GRAB_ATP_ fluorescence at 35 and 70 mmHg and with 50 mM KCl. Scale bar 20 µm. **B)** GRAB_ATP_ recordings of ATP release via pmCxL30 expressed in HeLa cells, shows that it forms an ATP permeable channel that is voltage sensitive and opened by CO_2_. Bottom recording, control recording from HeLa cells expressing only GRAB_ATP_, showing that there is no depolarisation or CO_2_ evoked ATP release in the absence of pmCxL30 expression. **C)** Summary data of depolarisation and CO_2_-evoked ATP release from 5 transfections. **D)** iGluSnFr recordings of glutamate release via pmCxL30 expressed in HeLa cells, shows that it forms a glutamate permeable channel that is voltage sensitive and opened by CO_2_. Bottom recording, control recording from HeLa cells expressing only iGluSnFr, showing that there is no depolarisation or CO_2_ evoked glutamate release in the absence of pmCxL30 expression. **E)** Summary data for the CO_2_-dependence of ATP and glutamate release from 5 transfections.

As many vertebrate connexins are permeable to ATP and glutamate, we co-expressed pmCxL30 with genetically encoded sensors for either ATP or glutamate to provide a real-time assay of channel opening. By analogy with vertebrate connexins, we used two physiological gating stimuli to trigger opening of putative pmCxL30 channels: membrane depolarisation induced by a high K^+^ stimulus and an increase in PCO_2_ from 20 to 70 mmHg. Both of these stimuli induced release of ATP or glutamate (Fig 2A-E). In HeLa cells that expressed only the genetically encoded sensors, neither a change in PCO_2_ nor KCl evoked analyte release. This showed that pmCxL30 was absolutely required for these responses.

We next investigated representatives from the CxL domains we identified in the polychaete *Owenia fusiformis* and the brachiopod *Lingula anatine* (Fig 3; Supplementary Fig 7). The polychaete molecules ofCxL26 and ofCxL40 were both opened by depolarisation but ofCxL26 was also sensitive to CO_2_ (Fig 3A). Meanwhile, the brachiopod CxL domain, laCxL27, was insensitive to CO_2_ but was opened by depolarisation (Fig 3B). We further sought to evaluate CO_2_ sensitivity in the connexins through analysis of the connexin genes of lamprey and Tunicates. Lamprey connexin genes map onto the major clades defined in the jawed vertebrate connexin molecular phylogeny, whereas the connexins of tunicates are largely distinct from vertebrate connexins. In both these early-branching chordates, we find evidence for sensitivity of these channels to depolarisation and CO_2_ (Fig 3C-D). Interestingly, the *Cnidarian* CxLs had a similar dose sensitivity to CO_2_ to the *Ciona* connexin, ciCx30.9, (compare Figs 3C and 2E) and other chordate connexins ^20, 22, 43^.

**Figure 3.**
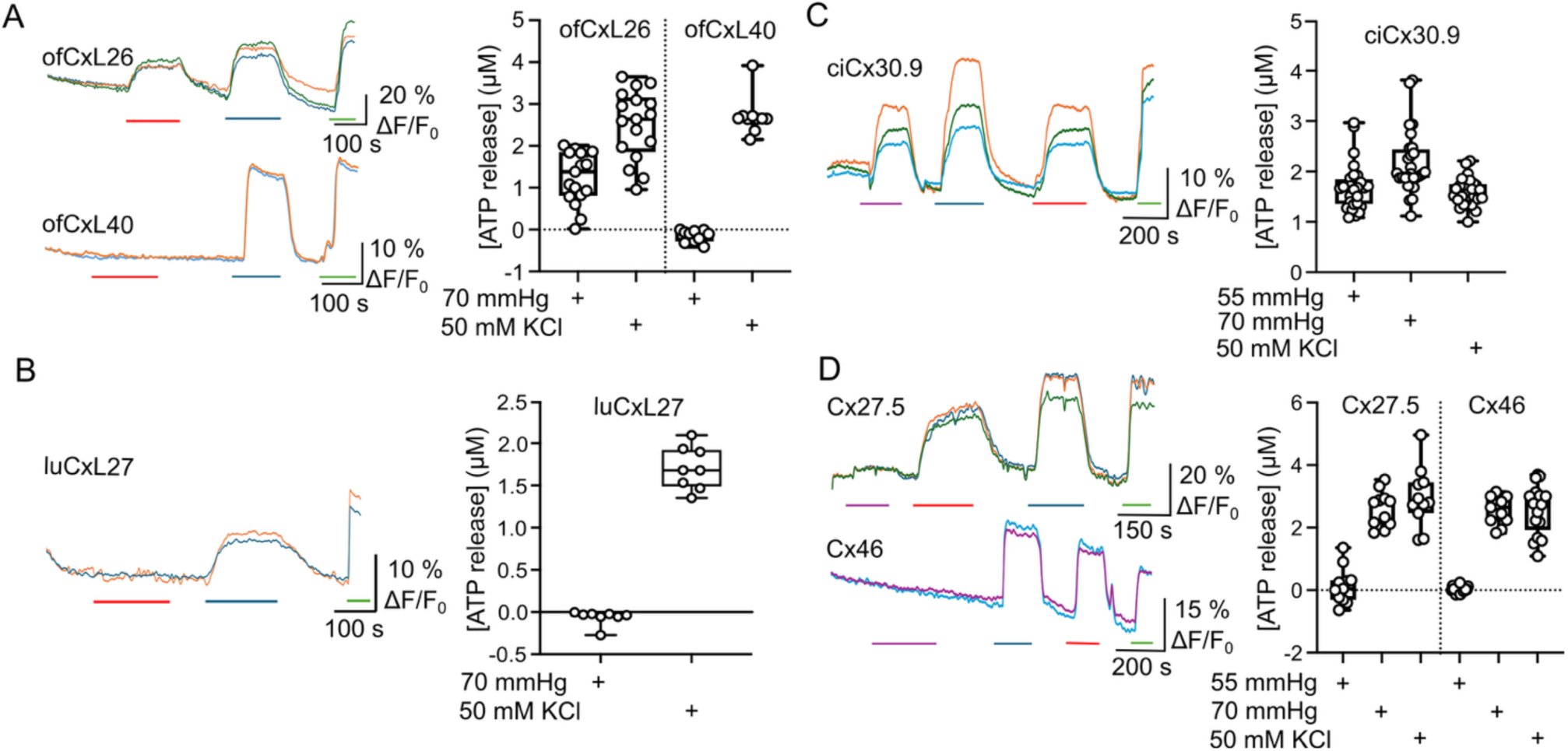
Connexin like molecules are present in annelid and brachiopod, and ancient chordates possess CO_2_ sensitive connexins. **A)** Two polychaete connexin-NLRs form membrane channels. ofCxL26 is sensitive to both membrane depolarisation (blue bar) and CO_2_ (red bar) whereas ofCxL40 is sensitive only to membrane depolarisation. **B)** luCxL27 from a brachiopod forms membrane channels that are voltage sensitive but not CO_2_ sensitive. **C)** A tunicate connexin, ciCx30.9, possesses the vertebrate carbamylation motif and is sensitive to membrane depolarisation and CO_2_. **D)** Two connexins from the lamprey (Cx27.5, an orthologue of Cx26, and Cx46) also possess the carbamylation motif and are sensitive to both membrane depolarisation and CO_2_. Purple bar indicates application of 55 mmHg PCO_2_ and red bar 70 mmHg PCO_2_. Data for all box plots were obtained from five transfections.

Our data show that these invertebrate CxL genes encode channels that share functional similarity to vertebrate connexins: they are gated by the same stimuli and form pores that are sufficiently large to permit passage of small molecules such as ATP and glutamate.

### Connexin-like domains predominantly form hemichannels

The canonical function of chordate connexins is to form gap junction channels that provide an aqueous passage for ions and small molecules between cells. Gap junction channels are dodecameric structures with the two hexameric hemichannels docking via a series of hydrogen bonds formed between the extracellular loops. Considering that there conflicting reports regarding the presence of gap junctions and connexin-mediated intracellular coupling in anthozoans ^44–48^, we investigated whether the CxL domains could form gap junctions (Fig 4). We transfected HeLa cells with either pmCxL30 (Fig 4A, F), edCxL50 (Fig 4B, F), and plCxL60 (Fig 4C, F) and looked for gap junction plaques between neighbouring expressing cells. These were not apparent and when we used a dye transfer assay to detect gap junction coupling, we found that the fluorescent dye, NBDG -a glucose derivative that readily passes through vertebrate connexin gap junction channels, did not transfer between cells. This indicates a lack of gap junction coupling for the three Cnidarian CxL molecules that we tested and likely stems from the CxLs possessing larger and more elaborate extracellular loops compared to the vertebrate connexins (Supplementary Fig 5).

**Figure 4.**
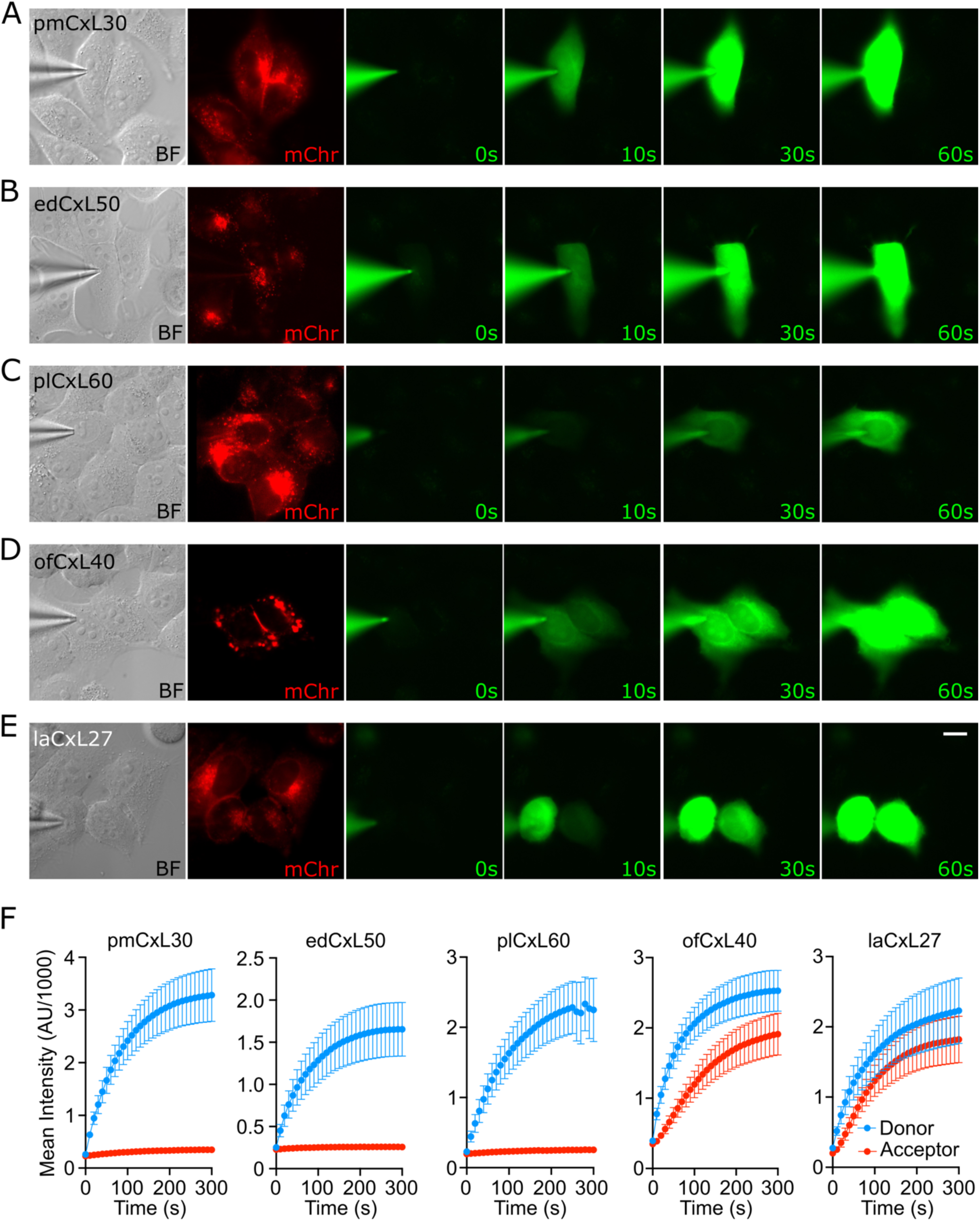
Gap junctions can be formed by protostome connexin-NLRs but have not been observed with cnidarian connexin-NLRs. A-E) Images of HeLa cells expressing various cnidarian and protostome connexin-NLRs. The left most image (BF) is a DIC image showing the patch electrode (filled with NBDG-glucose) recording from a single cell. The second image (mChr) from the left is mCherry emission showing expression of the mCherry tagged connexin-NLR. The remaining images show NBDG-glucose fluorescence. The numbers show the time after establishing the whole cell recording configuration. For the three cnidarian constructs (pmCxL30, edCxL50 and plCxL60, NBDG-glucose is confined to the recorded cell and does not enter any others. For the two protostome connexin-NLRs (ofCxL40 and laCxL27) the dye spreads to an adjacent cell within 10 s of establishing the whole cell configuration. Scale bar 15 µm. **F)** Measurements of fluorescence intensity for the donor cell (blue, recorded with patch electrode) and potential recipient cells (red). Whereas there is no permeation into the recipient cells with expression of the cnidarian constructs, the protostome constructs permit rapid permeation into the neighbouring recipient cells. N = 3 transfections for each construct.

We next tested the protostome CxL molecules. Cells expressing either ofCxL40 (polychaete derived) or laCxL27 (brachiopod derived) exhibited gap junction like plaques. We confirmed gap junction channel formation by both ofCxL40 and laCxL27 by observing the ready transfer of NBDG between coupled cells (Fig 4D-F).

### Fused domains do not interfere with connexin properties

We also selected homologues that possessed more extensive C-termini, including the NTP hydrolysing domain or NACHT domain. The anthozoan CxL-protein from the sea anemone *Exaiptasia diaphana* edCxL50 was expressed and although not CO_2_ sensitive, it was able to release both ATP and glutamate in response to a depolarising stimulus despite the presence of a NACHT domain (Fig 5A, B; Supplementary Fig 8). Both pmCxL62 (Fig 5C, D; Supplementary Fig 8) and plCxL60 (Fig 5E, F; Supplementary Fig 8) containing NACHT and caspase domains, respectively, exhibited CO_2_ sensitivity and voltage sensitivity.

**Figure 5.**
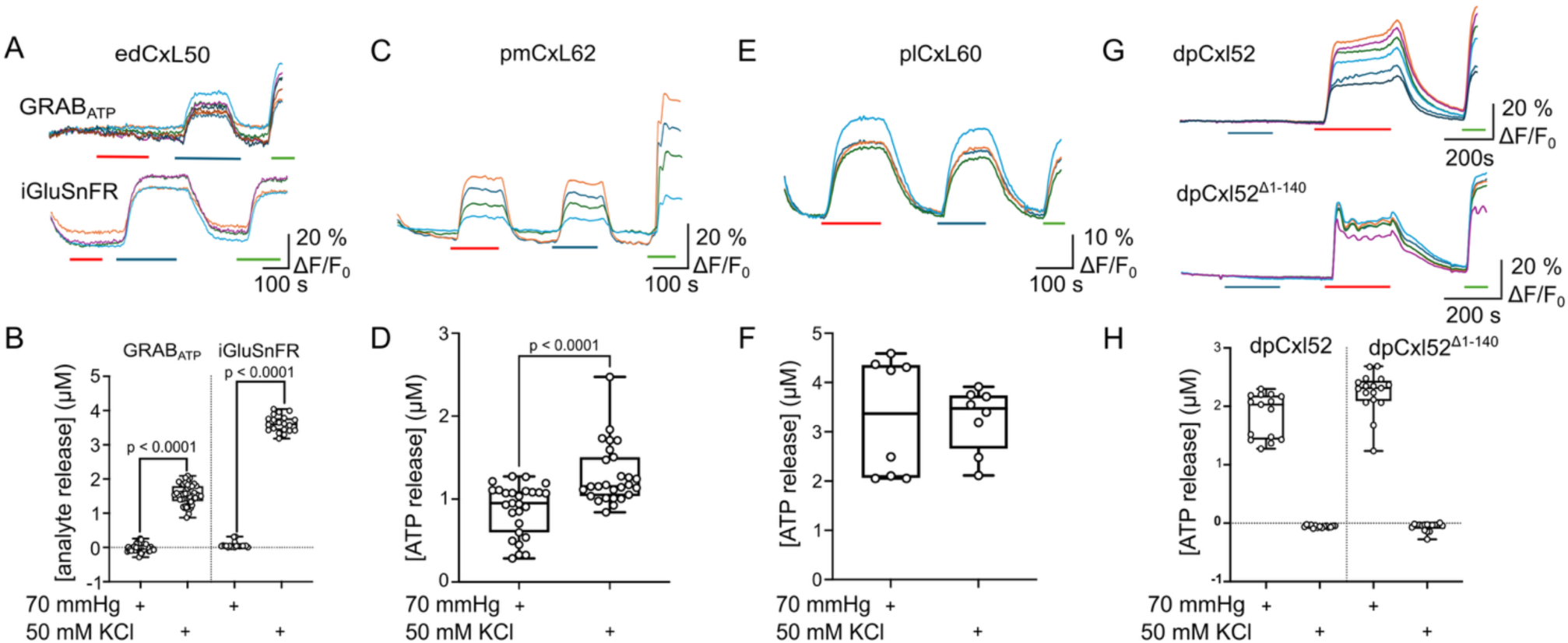
Cnidarian connexin-NLRs with multiple domains also form channels that can be voltage and CO_2_ sensitive. A, **B)** edCxL50 (NACHT domain) is insensitive to CO_2_ but can be opened by depolarisation and is permeable to both ATP and L-glutamate. **C, D)** pmCxL62 (NACHT domain) is sensitive to both membrane depolarisation and CO_2_. **E, F)** plCxL60 (caspase domain) is also sensitive to both membrane depolarisation and CO_2_. G, H) dpCxL52 (N-terminal TIR domain) is sensitive to CO_2_ but not membrane depolarisation. Deletion of the N-terminal TIR domain (dpCxL52^Δ1-140^) did not result in a gain of voltage sensitivity. **A, C, E, G)** Red bars 70mmHg, blue bars 50 mM KCl, green bars 3 µM calibrant. Data for all box plots were obtained from five transfections.

In our exploration of anthozoan connexin-NLRs we noticed some variants in which the genome annotation placed the connexin domain within the middle of the gene. One such example is dpCxL52 from *Desmophyllum pertusum*, which has a predicted TIR domain at the N-terminus, followed by the connexin domain and then the NLR. In a connexin, the N-terminus projects into the pore and is a key part of the gating mechanism. In dpCxL52 as this cannot be the case, we were curious as to whether it could form a channel, and if so how that channel could be gated.

We expressed dpCxL52 tagged with mCherry at the C-terminus in HeLa cells and observed punctate expression consistent with plasma membrane expression. We found that the channel could be opened by a PCO_2_ stimulus, but that it was insensitive to depolarisation as the high K^+^ stimulus did not evoke ATP release (Fig 5G, H; Supplementary Fig 8). It is interesting that dpCxL52 clearly had a gating mechanism, despite the lack of an N-terminal helix within the pore. The lack of voltage sensitivity might plausibly be explained by the presence of the TIR domain at the N-terminus effectively preventing the N-terminal helix from forming a gate within the pore that would typically be observed in a vertebrate connexin. We therefore removed the first 140 residues so that the protein started at M141 (dpCxL52^Δ1-140^). Interestingly the predicted structure had an N-terminal helix that projected into the pore. Nevertheless, while dpCxL52^Δ1-140^ could still be opened by changes in PCO_2_, it remained insensitive to depolarisation (Fig 5G, H; Supplementary Fig 8).

### CO_2_ sensitivity is a widespread feature of connexin-like domains

We recognised that some connexin-NLRs appeared to have a capacity for intersubunit carbamate bridge formation that we originally discovered in connexins ^21^ (Supplementary Fig 9). In some vertebrate connexins (e.g. exemplified by Cx26), there is a conserved carbamylation motif in which a Lys residue present in TM3 (K125) becomes carbamylated and interacts via a salt bridge with R104 in TM2 of the neighbouring subunit to mediate hemichannel opening ^21, 49–51^. A further residue in TM2, K108, becomes carbamylated and mediates CO_2_-dependent gap junction channel closure possibly via interaction with R216 in TM4 ^51^. In Cx43, the gap junction channels are insensitive to CO_2_, yet experience CO_2_-dependent hemichannel opening but the mechanism is somewhat more complex than that of Cx26. Cx43 hemichannel opening to CO_2_ still involves Lys residues equivalent to those identified in Cx26. In Cx43 there is potential for interaction between K144 in TM3 and either K105 or K109 in TM2 of the neighbouring subunit following carbamylation. Additionally, K105 and K109 of TM2 are sufficiently close to interact with K234 in TM4 of the neighbouring subunit. Mutational analysis implicates all four of these Lys residues in the CO_2_ sensitivity of Cx43 ^22^.

While the anthozoan and protostome CxLs did not possess the exact vertebrate carbamylation motif, we recognised that in pmCxL30 there is potential for a similar intersubunit interaction between residues K140 in TM3 and R232 in TM4, following hypothesized carbamylation of K140 (Fig 6A, Supplementary Fig 9). Note that in the predicted fold of pmCxL30, the interactions that we see in vertebrate connexins between the relevant residues TM2 with those of TM3 or TM4 of a neighbouring subunit are not plausible. While we observed a similar potential for intersubunit interactions following carbamylation in several other connexin-NLRs, it was absent in many other connexin-NLRs.

**Figure 6.**
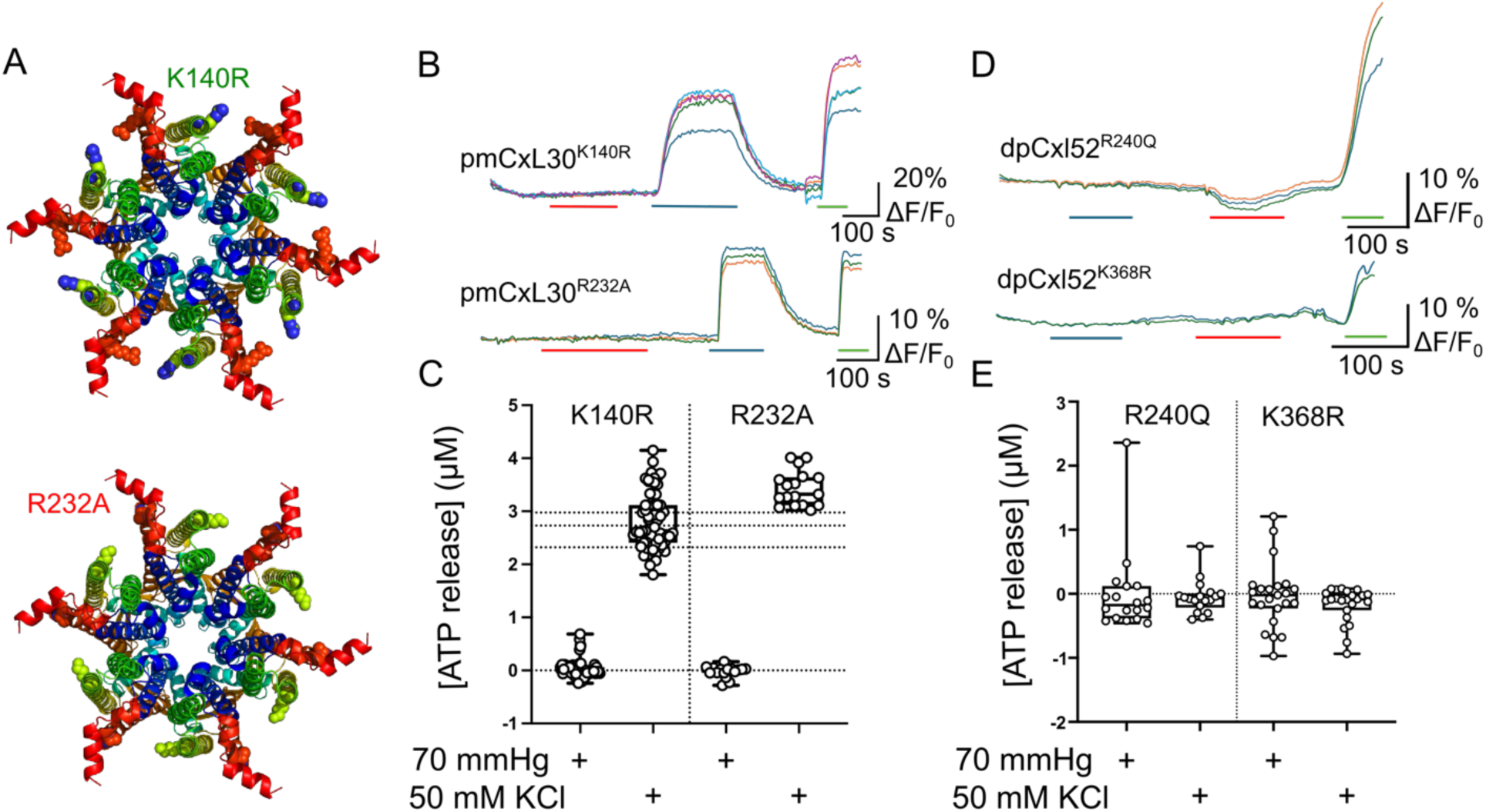
The CO_2_ sensitivity of connexin-NLRs is mediated via a carbamylation motif analogous to that of vertebrate connexins. **A)** The potential CO_2_ interacting residues in the predicted structure of pmCxL30. Mutations of K140R (in TM3) is predicted to prevent carbamylation and R232A (in TM4) to prevent a possible intersubunit salt bridge that could be formed following carbamylation of K140 and subsequent interaction with R232. **B, C)** The mutations K140R and R232A abolished CO_2_ sensitivity but did not alter voltage sensitivity. **D, E)** The CO_2_ sensitivity of dpCxL52 was also predicted to depend on intersubunit salt bridge formation following potential carbamylation of K368 (TM4) interacting with R240 (TM2). The mutations K368R and R240Q abolished CO_2_ sensitivity. Data for all box plots were obtained from five transfections.

We next tested whether the CO_2_ sensitivity of pmCxL30 depended upon the residues that we had identified in the AF3 predicted structures. To prevent carbamylation from occurring, we mutated Lys140 to Arg. This completely blocked sensitivity to CO_2_ but not sensitivity to depolarisation (Fig 6B, C; Supplementary Fig 10). To block interaction of the carbamylated Lys140 across the subunit boundary, we mutated Arg232 to Ala. This too abolished the CO_2_ sensitivity without affecting voltage sensitivity (Fig 6B, C; Supplementary Fig 10). Similarly, inspection of the predicted structure of dpCxL52 revealed a potential structural motif for carbamylation between R240 and K368 (Supplementary Table 7). Without the 140 amino acid TIR domain, these residues would be R100 and K228, which puts them in a similar position within the connexin-like structure to those residues in other CO_2_-sensitive connexin-NLRs, albeit forming the potential bridge in the reverse direction. The mutations R240Q and K368R individually abolished CO_2_ dependent opening of the channel (Fig 6D, E; Supplementary Fig 10). It would seem therefore that in dpCxL52, carbamylation occurs on a Lys residue in TM4 which then interacts with an Arg residue in TM2 (Supplementary Fig 9). The predicted structure of the polychaete molecule, ofCxL26, also suggested the presence of the motif with carbamylation of K108 potentially interacting with R197 of the neighbouring subunit (Supplementary Table 7). Mutation of the critical Lys residue to Arg (K108R) abolished CO_2_ sensitivity but did not affect voltage sensitivity (Supplementary Fig 11). Our hypothesis, that the CO_2_ sensitivity of the anthozoan and protostome connexin-NLRs depends upon the carbamylation motif, is further supported by our observation that further members of this protein family which lack the motif are insensitive to CO_2_ (edCxL50, ofCxL40, laCxL27 – Supplementary Table 7). It is interesting to note that the carbamylation motif in the early branching chordates is consistent with the anthozoan and protostome motif (Supplementary Tables 8, 9), yet the formation of the carbamate bridge occurs between TM3 and TM2, mirroring the vertebrate structural motif (Supplementary Table 7).

## Discussion

In our study, we reveal the presence of CxL domains in three metazoan phyla outside of Chordata, pushing the origins of these domains to at least the cnidarian-bilaterian ancestor 600-700 million years ago ^52^. CxL domains are distinct from innexins/pannexins and share sequence and structural similarity to chordate connexins. Our physiological analyses demonstrate that these domains form membrane-localised channels in human cell lines, are present as either gap junctions (protostomes) or hemichannels (anthozoans) and allow the passage of small molecules across membranes in response to depolarisation. Our work redefines the evolutionary history of the connexin gene family, demonstrates shared functional principles between invertebrate CxLs with vertebrate connexins, and provides important clues to the ancestral functional of these domains.

The CxL domains were identified by the similarity of their fold to vertebrate connexins. Our results show that they are functionally similar too. The predictions from AF3 suggest that they most plausibly form either hexamers or heptamers. That they are readily permeable to ATP and glutamate and must therefore form a pore large enough to accommodate these molecules, supports these predictions. They are also opened by two physiological stimuli that open connexin hemichannels: transmembrane depolarisation and an increase in PCO_2_. The voltage sensitivity can be at least partly understood by the predicted projection of a charged N-terminus into the pore. Similarly, the PCO_2_ sensitivity of some anthozoan and protostome CxL molecules appears to result from the same post-translational modification, carbamylation of a Lys residue, as vertebrate connexins.

Our observation that CO_2_-sensitivity among invertebrate CxLs further emphasises the importance of CO_2_-mediated gating in hemichannels. Sensing of CO_2_, a waste product of cellular metabolism, is distinct from pH sensing and its physiological importance in animals is increasingly being recognised^53^. In mammals, CO_2_-mediated gating of connexins is involved in respiratory responses to hypercapnia ^14, 54, 55^, and has been linked to conditions including X-linked Charcot-Marie-Tooth disease ^7^ and KID syndrome ^56^. While the carbamylation of a Lys residue is consistent between vertebrate connexins and anthozoans/protostomes CxLs, the nature of the interaction to a charged residue across the subunit boundary differs. In vertebrate connexins the interactions are between a carbamylated Lys in TM3 with a Lys/Arg in TM2. This exact pattern of interaction is not present in the anthozoan and protostome CxLs. Instead, most interactions take place between a carbamylated Lys residue in TM2 or TM3 with a Lys/Arg in TM4. Interestingly the CO_2_ dependent opening of Cx43 hemichannels could reflect an intermediate state as it involves Lys residues in TM2, TM3 and TM4^22^. The carbamylation motif in vertebrate connexins is highly conserved and comprises KF(R/H)FXG(where F corresponds to a hydrophobic residue and X to any residue). In invertebrates and ancient chordate connexins the consensus carbamylation motif is KFFFXF (Supplementary Table 8, 9). This similarity raises the possibility that the invertebrate carbamylation motif may be the evolutionary forerunner of the motif present in vertebrate connexins.

The presence of connexins in the cnidarian-bilaterian ancestor raises questions regarding the origins of these domains and the processes driving the diversification and composition of connexins and related channels (e.g., pannexins/innexins). The widespread prevalence and sequence diversity of CxLs in anthozoans and sporadic occurrence of connexin domains in non-chordate bilaterians is suggestive of widespread secondary gene loss. This is consistent with previous observations where early sequencing of anthozoans overturned the chordate-specific assignment of many genes that were missing from common lab models such as *Drosophila melanogaster* and *Caenorhabditis elegans* ^57–59^. It has been suggested that connexins arose in response to a bottleneck in innexins in early chordate evolution ^10^, which points to functional redundancy between the two families. However, the expansive repertoire of innexins in ctenophores^60^ and other cnidarians ^61, 62^ suggests that an innexin bottleneck may not have been responsible for the original emergence of connexins. The retention of CxLs in anthozoans alongside innexins likely stems from functional divergence between anthozoan CxLs and their innexin counterparts, mirroring vertebrate connexins and pannexins^63^ in that they are not mutually exclusive. Anthozoan CxLs presumably function in immune signalling as hemichannels, given their association with NLRs and that their elaborate extracellular loops likely prevent gap junction formation. In light of bilaterian non-gap junction forming exceptions such as mouse Cx23^64^ and Cx29 ^65^ (and the corresponding human ortholog Cx31.3 ^66, 67^), our findings suggest that hemichannel formation may have deeper evolutionary origins than previously recognised. As such, our findings underscore the need to reevaluate the long-standing assumption that the ancestral connexins formed gap junctions.

Connexins are a diverse family of proteins that are an essential component of healthy cells but have also been linked to pathophysiology and inflammation (e.g.^68–70^). Our screening revealed CxLs as N-terminal domains in NLRs, and alongside other immune-related domains (e.g. STING), in both anthozoans and protostomes suggestive of an ancestral role of CxLs in innate immunity. An ancestral innate immune role also helps contextualise the localisation of coral CxL-NLR expression to immune cell types ^33^ and our findings that anthozoan CxLs form hemichannels rather than gap junctions. Similar to pannexin1, the absence of gap junction formation would enable the release of signalling molecules into the extracellular space ^71^. Nevertheless, it is important to note that associations between connexins and innate immunity, particularly NLRs, are not limited to invertebrates. For example, connexin43 hemichannels prime and activate the NLRP3 inflammasome formation through ATP signalling ^70^ and this activity can be suppressed using hemichannel blockers ^72^ (e.g., Tonabersat). Further, certain connexins and NLRs are regulated by NF-κB ^73, 74^, and expressed in macrophages ^41, 75^. As such, our findings of putative ancestral links between CxLs and NLRs offers exciting new perspectives on vertebrate connexin function.

## Methods

### Bioinformatics and tree construction

CxL domains were identified through querying the Uniprot database for proteins possessing the IPR038359 (Connexin N-terminal domain superfamily) domain. The hits were filtered based on taxonomy to retain non-chordate proteins. Additional filters were applied to remove candidates from taxa whose CxL domains represented vertebrate contamination, were too short and/or whose predicted structures in AF3 were not consistent with known connexins (Supplementary Table 1).

Maximum likelihood trees were generated for CxL sequences from both chordate and non-chordates and for anthozoan CxLs. Sequences were aligned in MAFFT (v 7.487) ^76^ using the E-INS-I algorithm and a maximum likelihood tree was generated in IQTREE2 (v2.1.2) ^77^. The optimal substitution model was determined by ModelFinder ^78^. Trees were rooted using the minimal ancestor deviation (MAD) method ^79^ and visualised in iTOL (v7) ^80^.

To compare our CxL candidates to other membrane channels, we used EFI-EST ^81, 82^ to generate a sequence similarity network (SSN) for Uniprot domains belonging to the Connexin N-terminal domain superfamily (IPR038359), innexin (IPR000990), and pannexin domains (IPR039099) clustered at 50% (UniRef50). Filters were applied to retain hits with e-values less than 1-e^05^. Similarly, a connexin-only SSN was generated using the same e-value threshold but with the UniRef90 database. SSNs were visualised in Cytoscape. Further evaluation of remote homology CxLs and connexins was performed using HMM and structural approaches. Pairwise comparison of profile HMMs was performed in HHPred ^39^ using an HMM generated from alignment of anthozoan CxLs. Any anthozoan CxLs with a connexin PFAM (PF0029) annotation were excluded from the alignment. Foldseek ^38^ was used to compare structures predicted by AF3 for anthozoan CxLs and cnidarian innexins to predicted protein structures in the AFDB-PROTEOME.

As anthozoans were the most abundant non-chordate source of CxLs, we performed an additional screening for cnidarian CxLs. All RefSeq-annotated cnidarian genomes were downloaded from NCBI using NCBI Datasets ^83^. Isoforms were collapsed by taking the longest transcript and domains identified using a local InterProScan with the Cath3D and PFAM databases. Each proteome was evaluated for the presence of connexin, innexin, and pannexin domains using the aforementioned InterPro domain IDs. For proteins containing CxL domains, any additional domains were recorded. To evaluate the gene structure, the location of the CxL domain in each protein was identified using an alignment of all anthozoan CxLs and compared to coding exon boundaries in the RefSeq annotation.

### Solutions used

<u>Control (35 mmHg PCO_2_) aCSF:</u> 124 mM NaCl, 3 mM KCl, 2 mM CaCl_2_, 26 mM NaHCO_3_, 1.25 mM NaH_2_PO_4_, 1 mM MgSO_4_, 10 mM D-glucose saturated with 95% O_2_/5% CO_2_, pH 7.4.

<u>Hypercapnic (70 mmHg) aCSF:</u> 73 mM NaCl, 3 mM KCl, 2 mM CaCl_2_, 80 mM NaHCO_3_, 1.25 mM NaH_2_PO_4_, 1 mM MgSO_4_, 10 mM D-glucose, saturated with ∼12% CO_2_ (with the balance being O_2_) to give a pH of 7.4.

<u>Depolarising (35 mmHg PCO_2_) aCSF:</u> 77 mM NaCl, 50 mM KCl, 2 mM CaCl_2_, 26 mM NaHCO_3_, 1.25 mM NaH_2_PO_4_, 1 mM MgSO_4_, 10 mM D-glucose saturated with 95% O_2_/5% CO_2_, pH 7.4.

### Cell culture and transfection

The CxL gene sequence were synthesized by IDT and subcloned into the pCAG-GS-mCherry vector. DNA gBlock was amplified using PCR with primers (IDT). Plasmids were generated using Gibson assembly. Point mutations were introduced using Gibson assembly. Overlapping fragments both containing the desired mutation were PCR amplified with primers (IDT). The presence of the correct assembly or point mutation was confirmed by DNA sequencing (Plasmidasaurus). The CxL construct was inserted upstream of mCherry, with a short 12 amino acid linker (GVPRARDPPVAT).

pDisplay-GRAB_ATP1.0-IRES-mCherry-CAAX was a gift from Yulong Li (Addgene plasmid # 167582; http://n2t.net/addgene:167582; RRID:Addgene_167582)

pCMV(MinDis).iGluSnFR was a gift from Loren Looger (Addgene plasmid #41732; http://n2t.net/addgene:41732; RRID: Addgene_41732)

We used HeLa DH cells as an expression system, as they exhibit very low expression of endogenous connexins and have been used for this purpose by numerous authors over a long period. Parental HeLa DH cells (ECACC 96112022; Research Resource Identifier [RRID]: CVCL_2483) were grown in Low-glucose Dulbecco’s modified eagle medium (DMEM) (Merck Life Sciences UK Ltd; catalog no.: D6046) supplemented with 10% FBS (Labtech.com; catalog no.: FCS-SA) and 50μg/mL penicillin/streptomycin. The HeLa DH cells were plated onto coverslips at a density of 7.5 × 10^4^ cells per well of a 6 well plate and transiently transfected using a mixture of 1 μg each of the CxL construct and either the GRAB_ATP_ or iGluSnFR genetically encoded sensors and 3 μg PEI for 6hrs. Cells were imaged 48 hours after transfection. We used a protocol to measure ATP release from cells developed and described in our previous work (Butler and Dale, 2023).

### Live cell fluorescence imaging and analysis

Cells were transiently transfected with one pCAG-CxL-mCherry construct and one genetically encoded fluorescent sensor 48 h prior to imaging. Cells were perfused with control artificial cerebrospinal fluid (aCSF) until a stable baseline was reached before perfusion with either hypercapnic or high K^+^ aCSF. Once a stable baseline was reached after solution change, cells were again perfused with control aCSF, and when a stable baseline was reached, recordings were calibrated by direct application of 3 μM of the corresponding analyte. All cells were imaged by epifluorescence (Scientifica Slice Scope, Cairn Research OptoLED illumination, 60X water Olympus immersion objective, numerical aperture 1.0, Hamamatsu ImagEM EM-SSC camera; Metafluor software). Circularly permuted GFP (cpGFP) in the sensors was excited by a 470 nm LED, with emission captured between 504 and 543 nm. The CxL constructs had a C-terminal mCherry tag, which was excited by a 535 nm LED, and emission was captured between 570 and 640 nm. Only cells expressing both cpGFP and mCherry were selected for recording, with cpGFP images acquired every 4 s. For each condition, at least three independent transfections were performed with at least two coverslips per transfection. Analysis of all experiments was carried out in ImageJ (https://github.com/imagej/ImageJ) (Schneider et al., 2012). Cell recordings were corrected for any motion using the Image Stabilizer plugin (Li, 2008). Regions of interest (ROIs) were drawn around cells coexpressing both sensor and connexin. Median pixel intensity was plotted as normalized fluorescence change (ΔF/F_0_) versus time to give traces of fluorescence change. The amount of analyte release was quantified as concentration by normalizing to the ΔF/F_0_ caused by application of 3 μM of analyte, which was within the linear portion of the dose– response curve for each sensor. Release from a single cell was considered to be a statistical replicate.

### Patch clamp recording measurements of gap junction coupling

An MCI Cleverscope, Photometrics Prime camera, Cairn Instruments OptoLED illumination, and an Olympus 60x water immersion (NA 1.0) objective were used to visualize the cells under brightfield DIC, and mCherry expression via epifluorescence. Micromanager software was used to control the illumination and camera settings and to save images for offline analysis via ImageJ. Standard patch clamp techniques were used to make whole-cell patch clamp recordings from HeLa cells that expressed a given CxL construct as assessed by mCherry fluorescence. The intracellular fluid in the patch pipette contained: K-gluconate 130 mM, KCl 10 mM, EGTA 10 mM, CaCl_2_ 2 mM, HEPES 10 mM, pH adjusted to 7.3 with KOH and was adjusted with pure water to a final osmolarity of 295 mOsm. To detect gap junction coupling a fluorescent tracer, 2-Deoxy-2-[(7-nitro-2, 1, 3-benzoxadiazol-4-yl)amino]-D-glucose (NBDG), was dissolved at 200 µM in the patch recording fluid. Pairs of cells in close contact, both expressing mCherry were selected for recordings. Following the initiation of the whole-cell recording mode, images were collected every 10 s to assess the fluorescence intensity of NBDG (excited at 470 nM) in the donor and recipient cells. A single gap junction recording was considered to be an independent replicate.

## Acknowledgements

We would like to thank Phill Stansfeld for assistance with the structural prediction of the *N. vectensis* connexin-NLR hexamer.

