## Supplementary Material for "Deep evolutionary origins of the connexin gene family"

**Affiliations:**
[1] School of Life Sciences, University of Warwick, Coventry, CV4 7AL

| Species | Phylum | Class | N. connexin | N. innexin /pannexin |
|---|---|---|---|---|
| *Acropora digitifera* | Cnidaria | Hexacorallia | 28 | 1 |
| *Acropora millepora* | Cnidaria | Hexacorallia | 100 | 0 |
| *Acropora muricata* | Cnidaria | Hexacorallia | 46 | 1 |
| *Actinia tenebrosa* | Cnidaria | Hexacorallia | 4 | 1 |
| *Dendronephthya gigantea* | Cnidaria | Octocorallia | 33 | 0 |
| *Exaiptasia diaphana* | Cnidaria | Hexacorallia | 12 | 1 |
| *Montipora capricornis* | Cnidaria | Hexacorallia | 21 | 1 |
| *Montipora foliosa* | Cnidaria | Hexacorallia | 18 | 0 |
| *Nematostella vectensis* | Cnidaria | Hexacorallia | 10 | 1 |
| *Orbicella faveolata* | Cnidaria | Hexacorallia | 7 | 1 |
| *Pocillopora damicornis* | Cnidaria | Hexacorallia | 13 | 1 |
| *Pocillopora verrucosa* | Cnidaria | Hexacorallia | 54 | 1 |
| *Stylophora pistillata* | Cnidaria | Hexacorallia | 22 | 1 |
| *Henneguya salminicola* | Cnidaria | Myxosporea | 0 | 0 |
| *Myxobolus squamalis* | Cnidaria | Myxosporea | 0 | 0 |
| *Thelohanellus kitauei* | Cnidaria | Myxosporea | 0 | 0 |
| *Clytia hemisphaerica* | Cnidaria | Hydrozoa | 0 | 41 |
| *Hydra vulgaris T2T AEPx105* | Cnidaria | Hydrozoa | 0 | 16 |
| *Hydra vulgaris T2TAEP* | Cnidaria | Hydrozoa | 0 | 16 |
| *Hydractinia symbiolongicarpus* | Cnidaria | Hydrozoa | 0 | 27 |
| *Rhopilema esculentum* | Cnidaria | Scyphozoa | 1 | 0 |
| *Owenia fusiformis* | Annelida | Polychaeta | 10 | 23 |
| *Lingula anatina* | Brachiopoda | Lingulata | 1 | 27 |

**Supplementary Table 5. Abundance of genes with connexin (IPR038359) and innexin/pannexin (IPR000990) domains in cnidarians and connexin-hosting protostomes.**

| Sequence ID | Length | CxL Domain Length | N. CxL Coding Exons | Total Coding Exons | Other Domains |
|---|---|---|---|---|---|
| CAH1778566.1 | 983 | 268 | 4 | 5 | P-loop_NTPase; TPR-like_helical_dom_sf |
| CAH1779526.1 | 351 | 226 | 1 | 5 | RHIM |
| CAH1791861.1* | 218 | 201 | 1 | 1 | N/A |
| CAH1797686.1 | 1065 | 229 | 2 | 3 | P-loop_NTPase; TPR-like_helical_dom_sf |
| CAH1797688.1 | 947 | 259 | 4 | 4 | N/A |
| CAH1797689.1* | 223 | 213 | 1 | 1 | N/A |
| CAH1797845.1 | 456 | 229 | 2 | 6 | STING |
| CAH1797848.1 | 1111 | 228 | 2 | 3 | P-loop NTPase; TPR-like_helical_dom_sf |
| CAH1798086.1 | 405 | 192 | 4 | 7 | STING |
| CAH1798091.1 | 1099 | 223 | 2 | 3 | P-loop_NTPase; TPR-like_helical_dom_sf |

**Supplementary Table 6. Gene and protein structure of *Owenia fusiformis* CxLs.** Exon analyses were limited to only coding exons. * *indicates partial sequence.*

| Species | Accession | NACHT | LRR | Other | Motif | Loc | Phylum |
|---|---|---|---|---|---|---|---|
| *Pocillopora meandrina* | A0A3M6TUS9_POCDA pmCxL30 | No | No | No | *K140, R232* | TM3, TM4 | Cnidaria |
| *Pocillopora meandrina* | A0AAU9X471_9CNID pmCxL62 | Yes | No | No | K130, R227 | TM3, TM4 | Cnidaria |
| *Porites lobata* | CAH3037244.1 plCxL60 | No | No | Caspase | K116-K212 | TM3, TM4 | Cnidaria |
| *Desmophyllum pertusum* | A0A9W9Y920_9CNID dpCxL52 | No | No | TIR | *R240, K368* | TM2, TM4 | Cnidaria |
| *Pocillopora damicornis* | XP_027053751.1* | No | No | No | K103, K220 | TM2, TM4 | Cnidaria |
| *Acropora digitifera* | XP_015779395.1* | No | No | No | K105, K214 | TM2, TM4 | Cnidaria |
| *Dendronephthya gigantea* | XP_028416259.1* | Yes | No | No | R111, K286 | TM2, TM4 | Cnidaria |
| *Acropora millepora* | XP_044167617.1* | Yes | Yes | PPP1R42 | K109, K244 | TM2, TM4 | Cnidaria |
| *Acropora millepora* | XP_044176766.1* | Yes | No | PPP1R42 | R114, K238 | TM2, TM4 | Cnidaria |
| *Nematostella vectensis* | XP_048579696.1* | Yes | Yes | RNA1 | R106, K248 | TM2, TM4 | Cnidaria |
| *Pocillopera verrucosa* | XP_066025853.1* | Yes | Yes | No | K133, R257 or K261 | TM3, TM4 | Cnidaria |
| *Pocillopera verrucosa* | XP_066028316.1* | Yes | Yes | RNA1 | R107, K245 | TM2, TM4 | Cnidaria |
| *Acropora muricata* | XP_067022978.1* | Yes | Yes | PPP1R42 LRR_RI | R112, K240 | TM2, TM4 | Cnidaria |
| *Owenia fusiformis* | A0A8J1U2R9_OWEFU ofCxL26 | No | No | No | *K108,* R197 | TM2, TM4 | Annelid |
| *Ciona intestinalis* | F6T0Z3 ciCx30.9 | No | No | No | K128, K104 | TM3 TM2 | Chordate (Tunicata) |
| *Petromyzon marinus* | S4RXA7 Cx27.5 | No | No | No | K120, R104 | TM3 TM2 | Chordate (Agnatha) |
| *Petromyzon marinus* | S4RXA6 Cx46 | No | No | No | K152, R101 | TM3 TM2 | Chordate (Agnatha) |
| *Homo sapiens* | NP_003995.2 Cx26 | No | No | No | *K125, R104* | TM3, TM2 | Chordate (Mammalia) |
| *Homo sapiens* | NP_001103689.1 Cx30 | No | No | No | K125, R104 | TM3, TM2 | Chordate (Mammalia) |
| *Homo sapiens* | NP_000157.1 Cx32 | No | No | No | *K124, K104* | TM3, TM2 | Chordate (Mammalia) |
| *Homo sapiens* | NP_000156.1 Cx43 | No | No | UBQLN4 | *K234, K144, K109, K105* | TM4, TM3, TM2, TM2 | Chordate (Mammalia) |
| *Homo sapiens* | NP_005258.2 Cx50 | No | No | No | *K140, K105* | TM3, TM2 | Chordate (Mammalia) |

**Supplementary Table 7. Incidence of $CO_2$ sensitivity and location of the carbamylation motif in Cnidarian CxL-NLRs and chordate connexins.** Role of italicized residues has been tested by mutagenesis. * Indicates that $CO_2$ sensitivity is predicted but not experimentally tested.

| | | |
|---|---|---|
| A0A3M6TUS9_POCDA/134-164 | **KLFLAY** | Cnidarian |
| A0AAU9X471_9CNID/124-154 | **KLFIAY** | Cnidarian |
| CAH3037244.1/111-144 | **KHLKTF** | Cnidarian |
| A0A8J1U2R9_OWEFU/108-141 | **KEMSPK** | Polychaete |
| F6T0Z3_CIOIN/128-134 | **KAAFSQ** | Tunicate |
| F6QCF4_CIOIN/150-156 | **KVPITK** | Tunicate |
| F6UBJ9_CIOIN/175-182 | **KLRRTK** | Tunicate |
| S4RXA7_PETMA/120-125 | **KPPIDG** | Lamprey |
| S4RXA6_PETMA/152-157 | **KVCIKG** | lamprey |

**Supplementary Table 8. Comparison of potential carbamylation motif in cnidarina, protostome and ancient chordate groups.** The first Lys group is proposed to be carbamylated and the next 5 residues orient appropriately to a Lys or Arg of the neighbouring subunit.

| | | | | | | | |
|---|---|---|---|---|---|---|---|
| Vertebrate | Sequence | **K** | **h** | + | **h** | **X** | **G** |
| | Conservation | 1.00 | 0.89 | 0.83 | 0.94 | | 1.00 |
| Cnidarian, protostome ancient chordate | Sequence | **K** | **h** | **h** | **h** | **polar or charged** | **h** |
| | Conservation | 1.00 | 0.78 | 0.67 | 0.67 | 0.67 | 0.56 |

**Supplementary Table 9. Comparison of consensus vertebrate carbamylation motif with the consensus anthozoan, protostome, and ancient chordate motif.** h -hydrophobic residue, + positively charged residue.

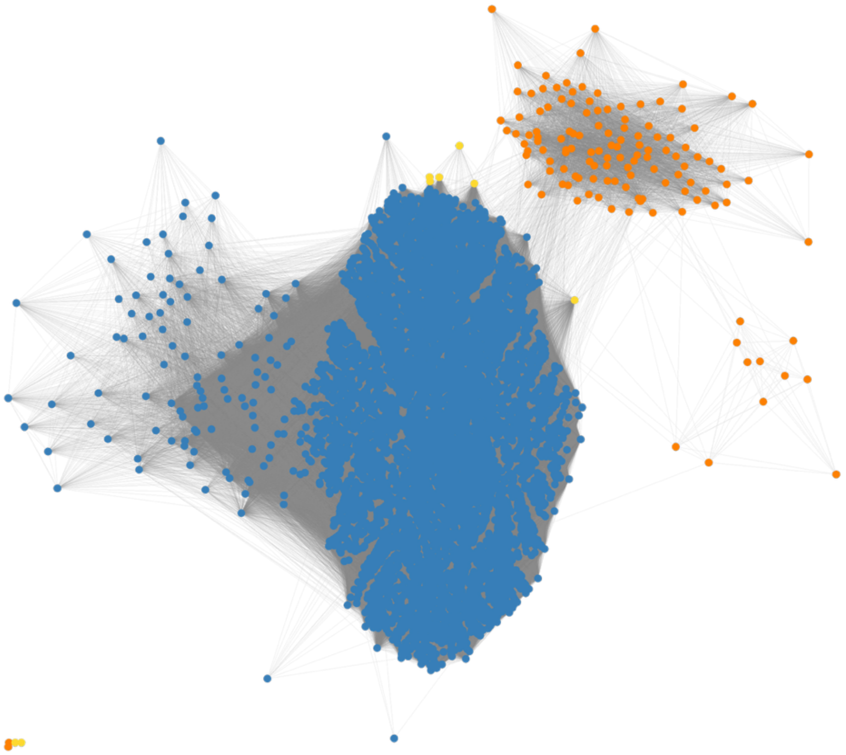

**Supplementary Figure 1: Anthozoan connexin-like domains share significant sequence similarity with chordate connexins.** Clustering network of protein domains annotated as members of the connexin (IPR038359) family in the UniRef90 database. Nodes are coloured according to taxonomy (Invertebrates excluding cnidarians and tunicates – yellow; cnidarians – orange; chordates – dark blue). Edges connect nodes sharing significant similarity (e-value < $1e^{-5}$).

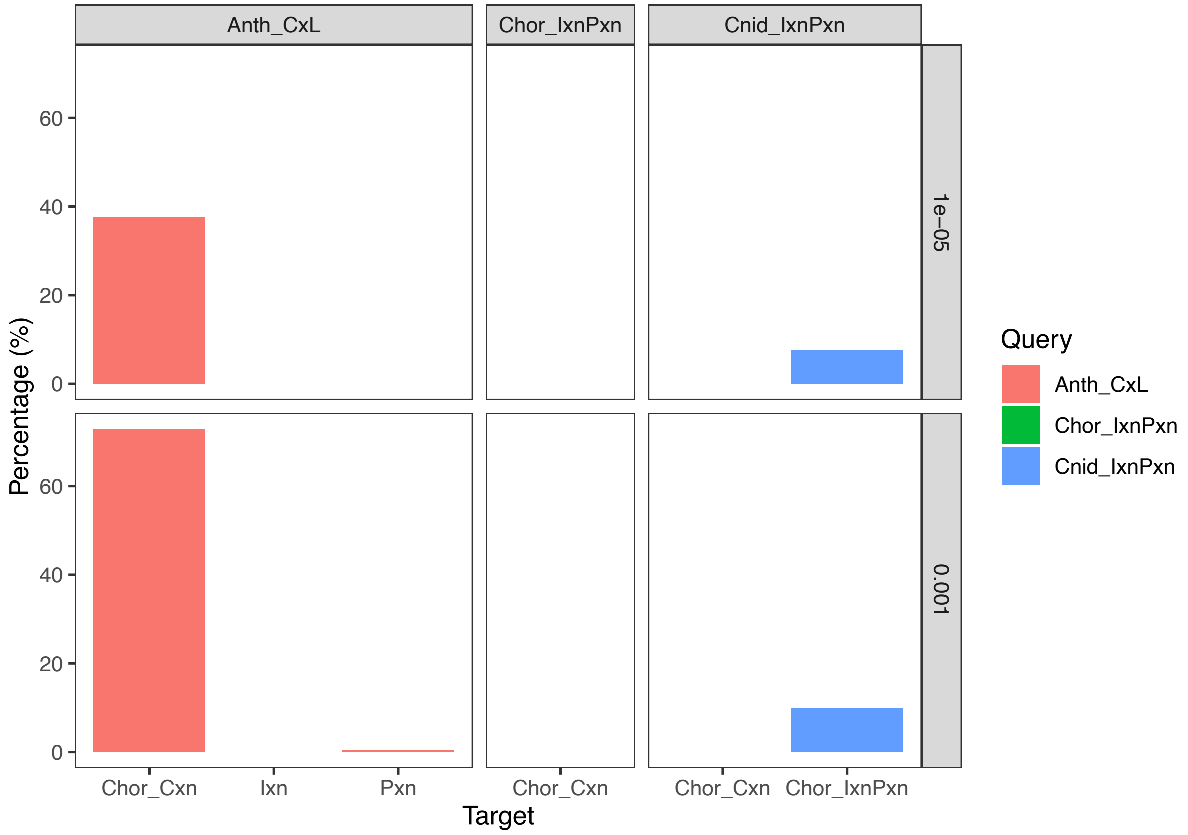


**Supplementary Figure 2. Anthozoan CxL-like domains consistently retrieve BLAST hits from chordate connexins and at a comparable rate to known homologous genes.** Bars represented the percentage of query domains that successfully match different targets across two E-value thresholds ($1E^{-3}$ and $1E^{-5}$). While cnidarian innexins/pannexins return proportionally fewer matches, these BLAST hits are composed of 78% and 100% of the anthozoan innexins/pannexins (n=9) at E-value thresholds of $1E^{-3}$ and $1E^{-5}$, respectively.

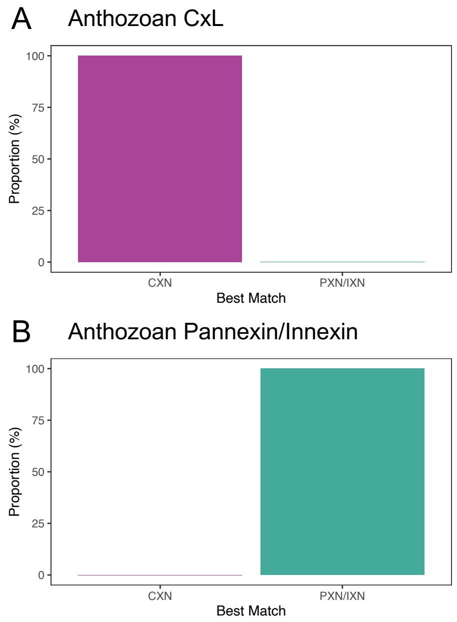


**Supplementary Figure 3. Anthozoan connexin-like and pannexin/innexin domains best match their members of their respective gene families based on structural comparisons.** Bars show the relative proportion of best hits belonging to each gene family for anthozoan connexin-like (A; n=30) and anthozoan pannexins/innexins (B; n=9).

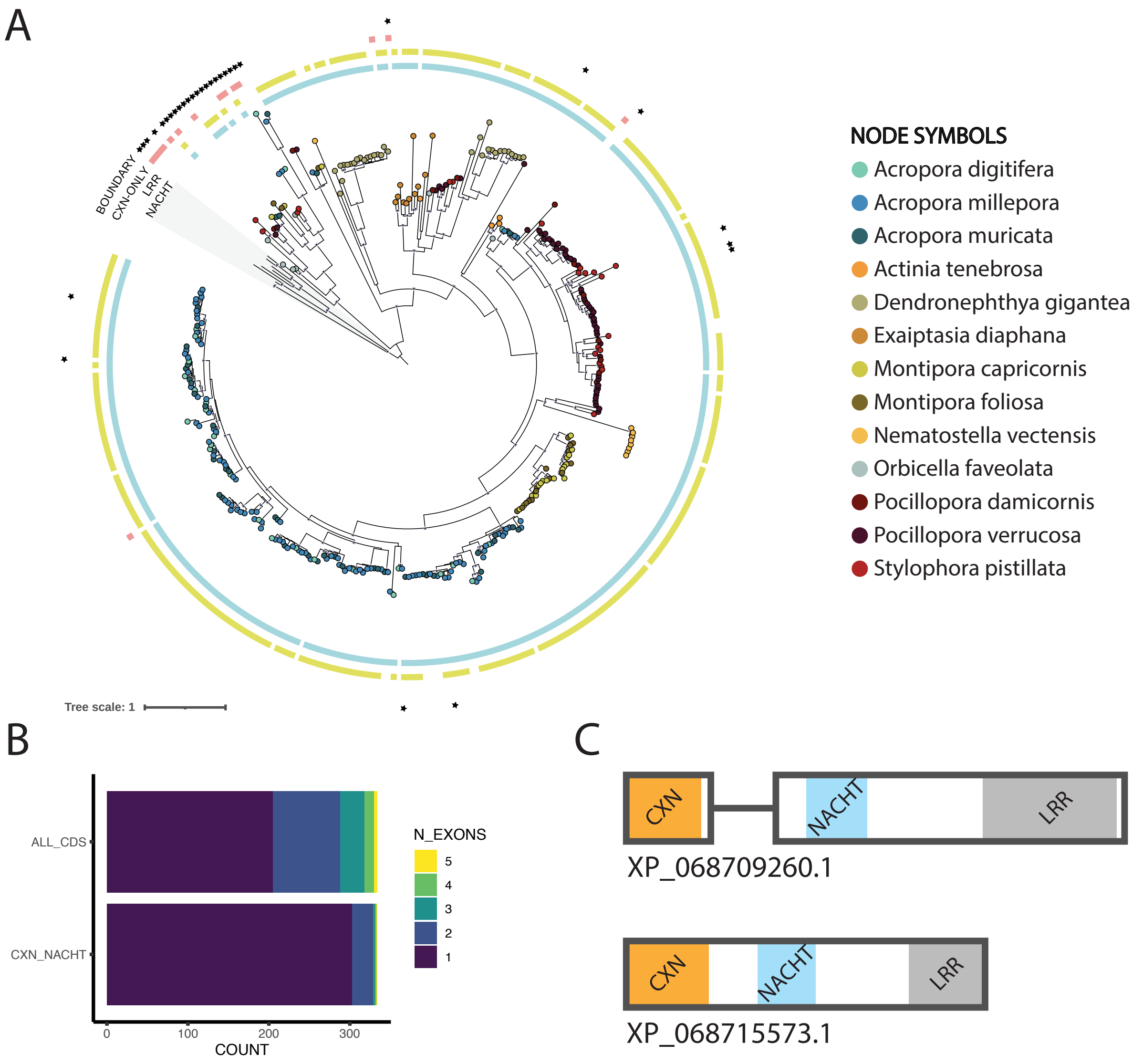


**Supplementary Figure 4. Cnidarian CxL-domains are a diverse family that are found alongside NACHT and LRR domains, with exceptions correlating with altered gene structures and largely restricted to a single clade.** A) Maximum likelihood tree with the presence of NACHT and LRR domains labelled as external squares. Proteins composed of only the connexin domain are labelled with red squares. Node symbols are coloured according to species and stars indicate proteins with an exon-intron boundary within 20 amino acids of the connexin domain. B) Number of exons within the span of the CDS (top) or within the range of the connexin and NACHT domains (bottom). C) Gene structure within the CDS for example genes with and without an exon-intron boundary located adjacent to the connexin domain.

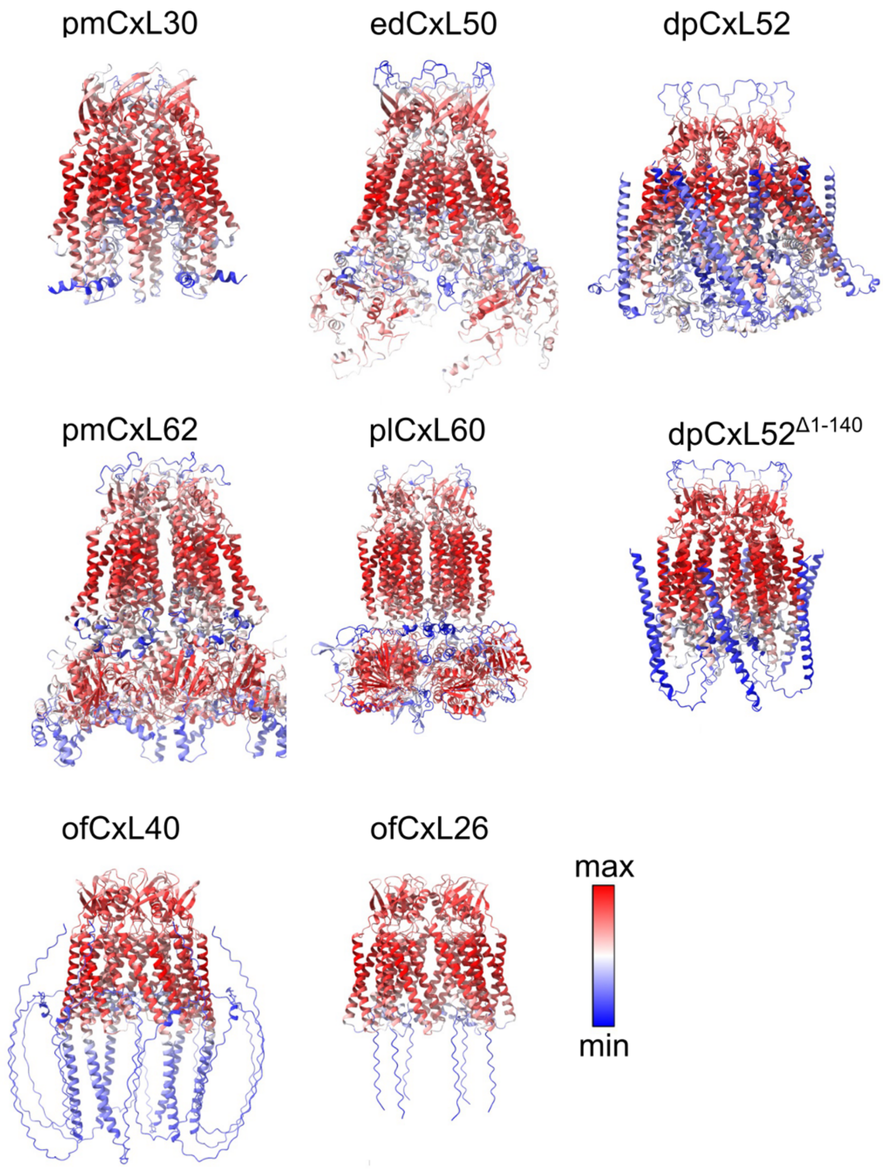


**Supplementary Figure 5. Alphafold3 predicted structures for the genes studied.** They are shown as hexamers and coloured according to the confidence

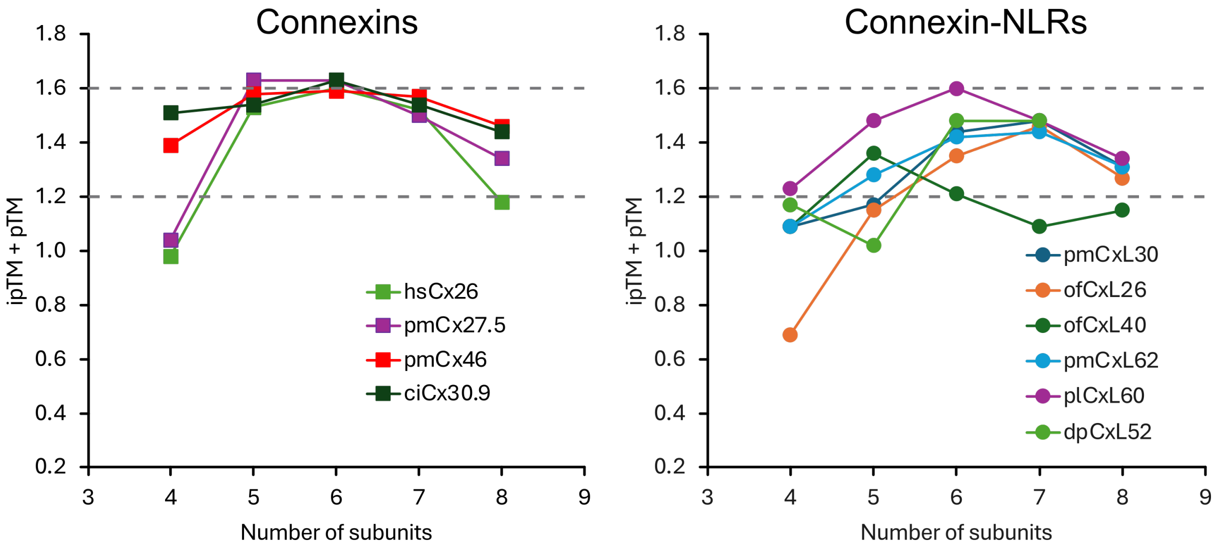


**Supplementary Figure 6. Confidence for predicted structures assembled with different numbers of Connexin and CxL subunits as measured by the sum of ipTM and pTM.** The dashed grey lines indicate the confidence grey zone in which a structure may be plausible. Below 1.2 the confidence is very low, above 1.6 it is very high. AF3 correctly predicts hexamers as the most plausible structures for connexins. Most CxL molecules gave the highest confidence scores when assembled as hexamers or heptamers. hsCx26 -human Cx26, the other names as described in the text.

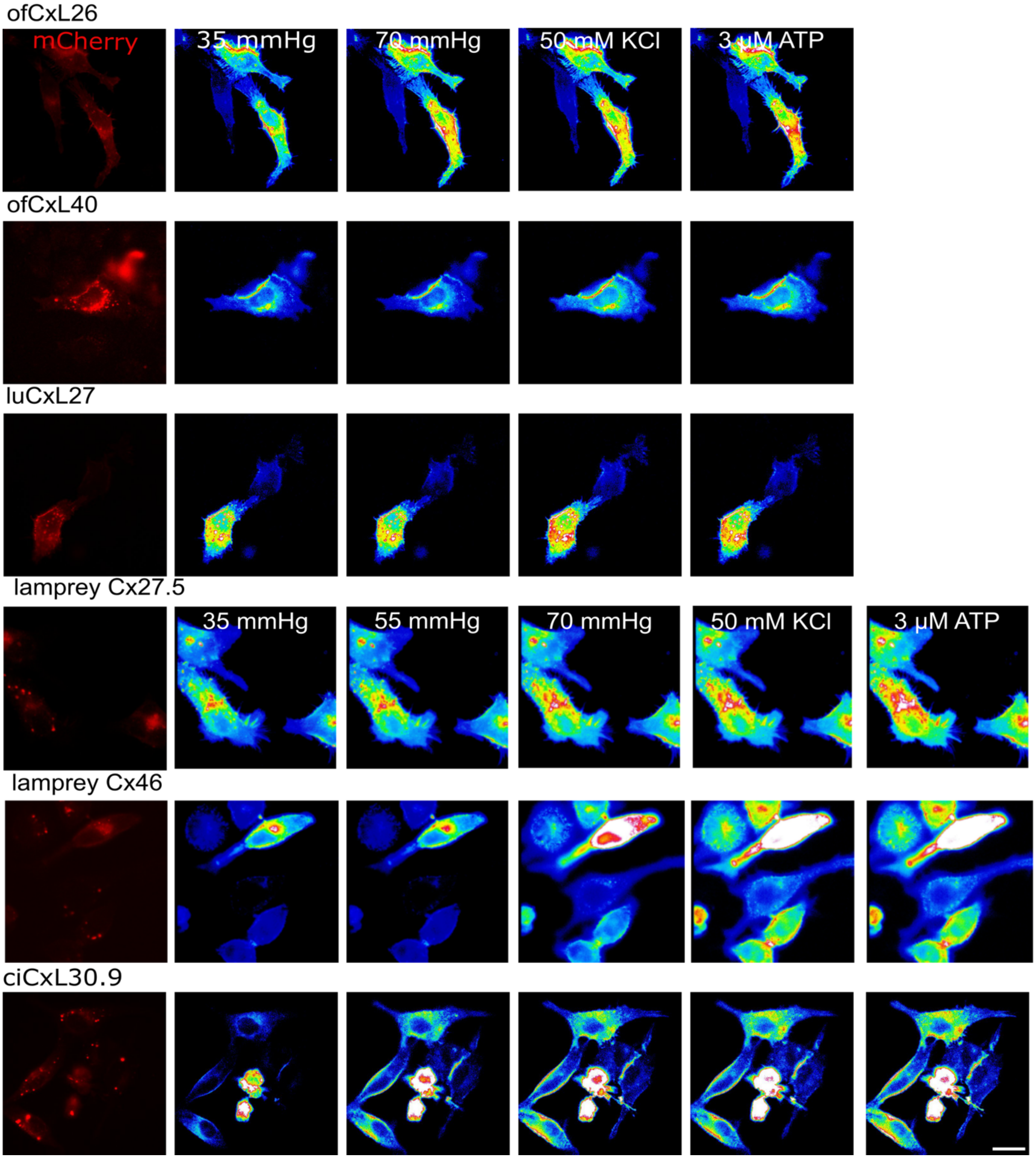


**Supplementary Figure 7. Representative images of the CxL or connexin expression pattern (mCherry) and GRAB$_{ATP}$ fluorescence.** The GRAB$_{ATP}$ fluorescence is presented as a pseudocolour image with blue being low and red being high. These images accompany Figure 3. Scale bar 20 µm.

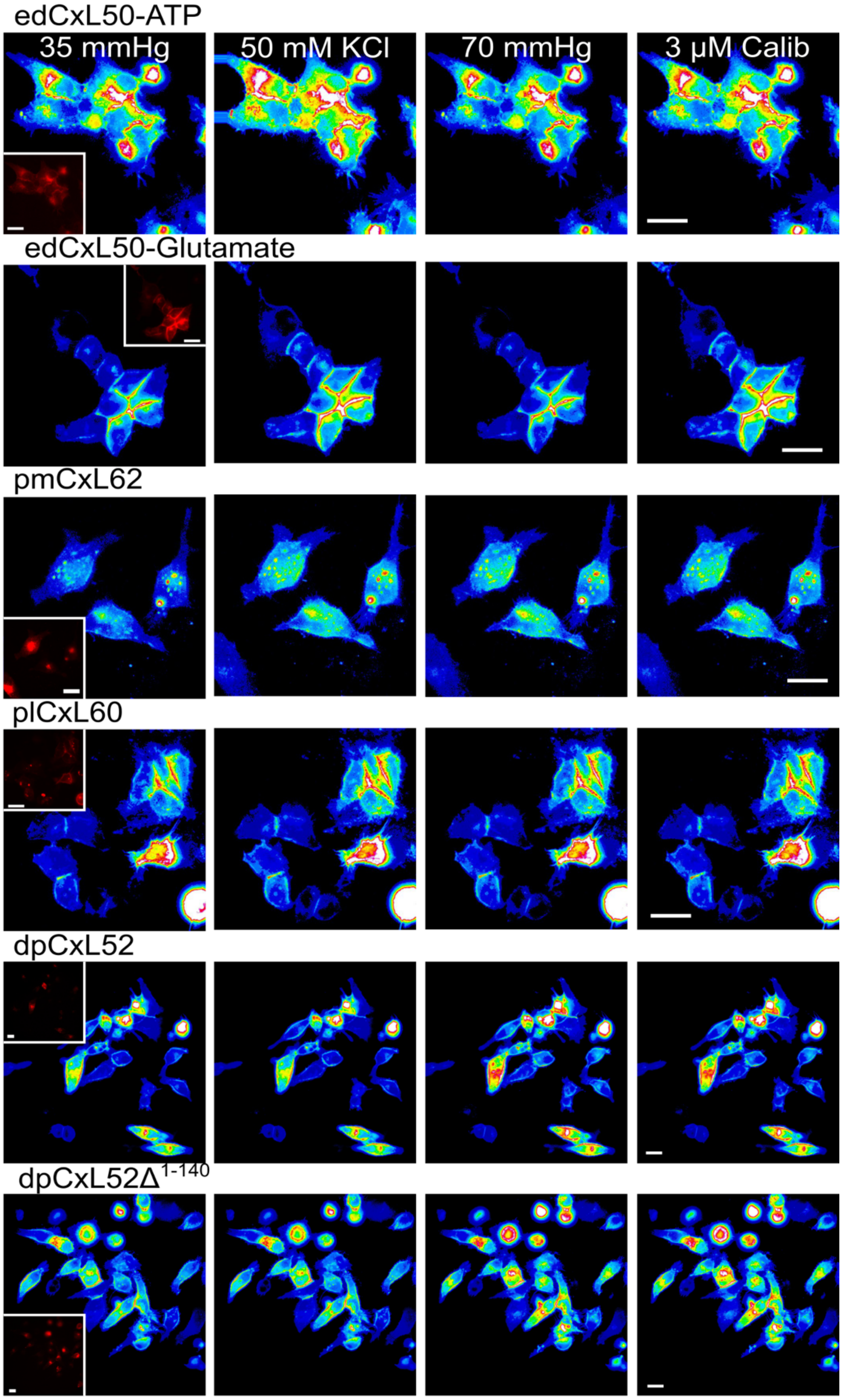


**Supplementary Figure 8. Representative images of the CxL expression pattern (mCherry) and GRAB$_{ATP}$ fluorescence.** The GRAB$_{ATP}$ fluorescence is presented as a pseudocolour image with blue being low and red being high. These images accompany Figure 5. Scale bar 20 μm.

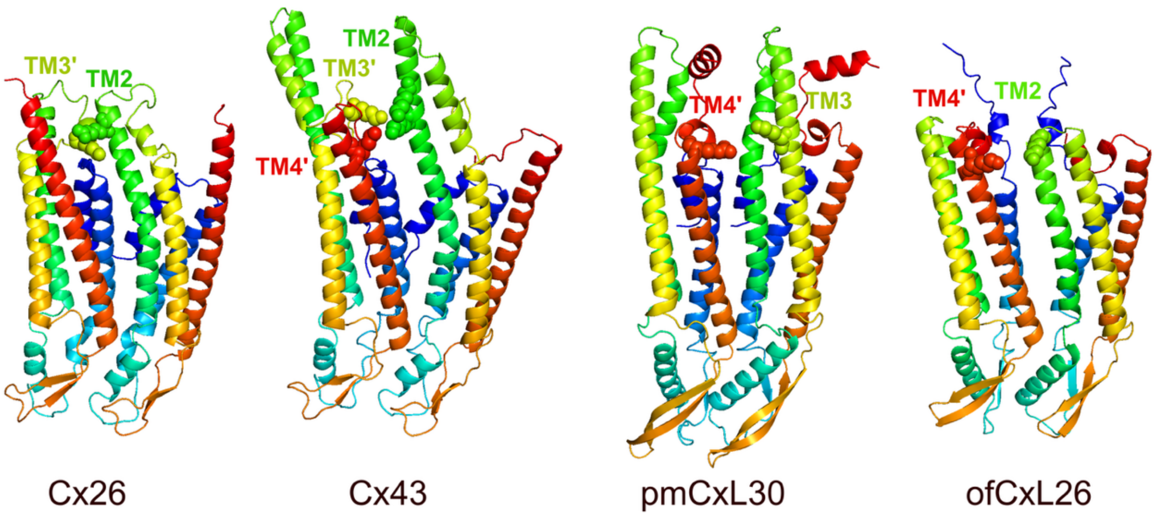


**Supplementary Figure 9.** Carbamylation motif in connexin and CxL domains. Ribbon diagrams represent the structures predicted by AlphaFold3 and putative amino acids involved in carbamylation motifs are shown with a space-filling representation.

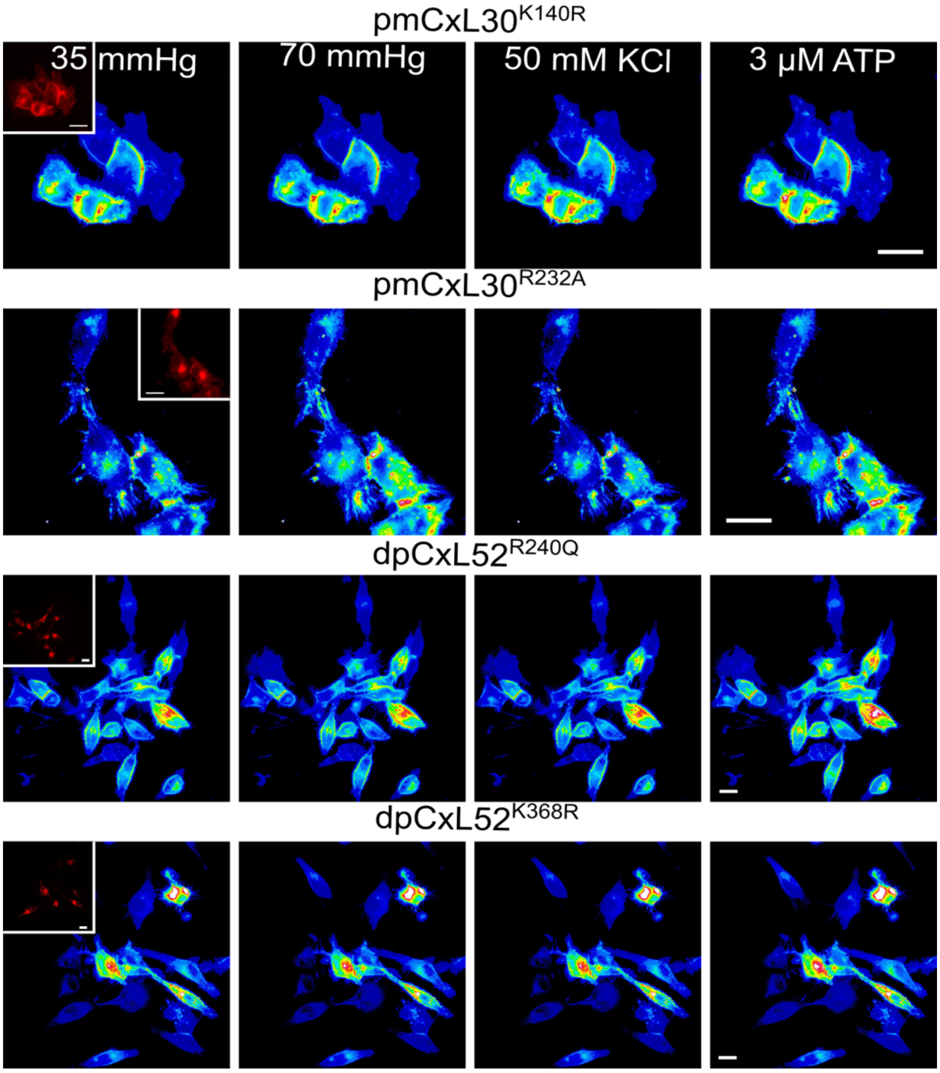


**Supplementary Figure 10. Representative images of the mutant CxL expression pattern (mCherry) and $GRAB_{ATP}$ fluorescence for pmCxL30 and dpCxL52.** In pmCxL30, K140R and R232A abolish $CO_2$ induced changes in $GRAB_{ATP}$ fluorescence, but do not affect depolarisation evoked ATP release. For dpCxL52, the mutations R240Q and K368R abolish $CO_2$ dependent ATP release. dpCxL52 is insensitive to transmembrane voltage. The $GRAB_{ATP}$ fluorescence is presented as a pseudocolour image with blue being low and red being high. These images accompany Figure 6. Scale bar 20 µm.

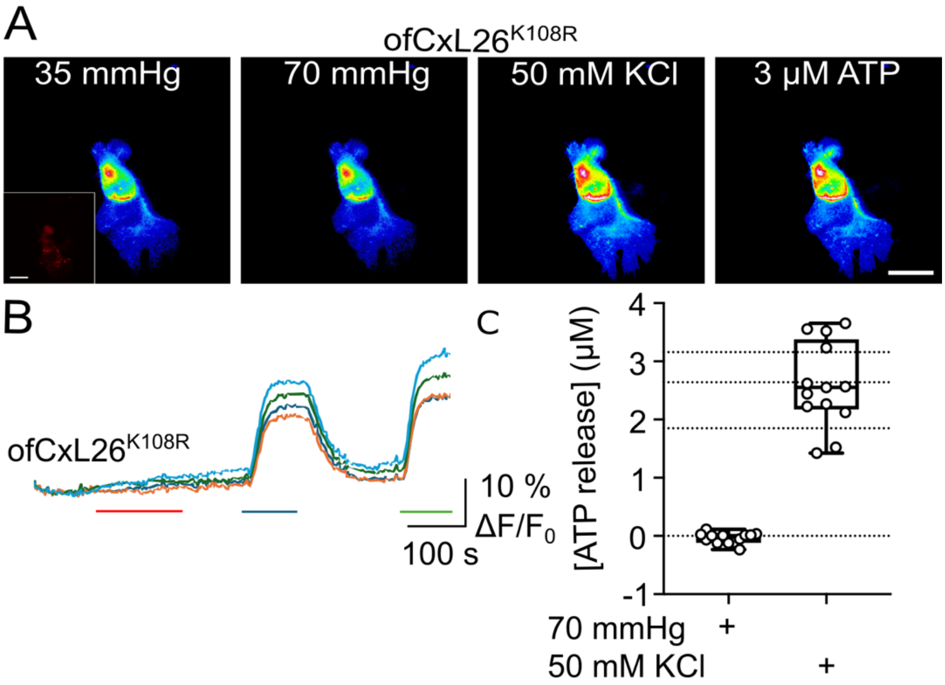


**Supplementary Figure 11. $CO_2$ dependent opening of ofCxL26 depends on putative carbamylation of K108. A)** Predicted structure and position of the interacting residues K108 and R197. **B)** Images of $GRAB_{ATP}$ fluorescence show that 70 mmHg $PCO_2$ does not evoke ATP release, whereas depolarisation with KCl does. **C)** Traces of $GRAB_{ATP}$ fluorescence showing the lack of response to 70 mmHg $PCO_2$ (red bar), the response to 50 mM KCl (blue bar) and 3 µM ATP calibration (green bar). **C)** Summary data from 5 independent transfections.
